# Genomic Epidemiology of Monkeypox Virus During the 2022 Outbreak in New York City

**DOI:** 10.1101/2024.07.23.604793

**Authors:** Saymon Akhter, Michelle Su, Jade C. Wang, Helly Amin, Faten Taki, Nelson De La Cruz, Moinuddin Chowdhury, Tyler Clabby, Erik Kopping, Victoria E. Ruiz, Mindy Leelawong, Julia Latash, Kimberly Johnson, Jennifer Baumgartner, Marcia Wong, Aaron Olsen, Randal C. Fowler, Jonathan E. Pekar, Jennifer L. Havens, Tetyana I. Vasylyeva, Joel O. Wertheim, Scott Hughes, Enoma Omoregie

## Abstract

New York City (NYC) was one of the hotspots of monkeypox virus (MPXV) infections in the United States during the multi-country mpox outbreak in 2022. This study used the most comprehensive dataset to date to investigate the genomic characteristics of MPXV in NYC. We performed Nextclade lineage assignment, phylogenetic and mutational analyses on 1,138 specimens from 748 individuals at the NYC Public Health Lab in the context of 2,968 MPXV sequences sampled globally. Nextclade lineage assignment showed B.1.12 as a NYC specific lineage and phylogenetic analysis showed NYC and North America specific clades. The majority of mutations showed signatures of APOBEC3 activity. When looking at the intra-host genomic diversity for MPXV, distinct MPXV genomic profiles were observed in 6.4% of individuals with multiple sampled specimens, with at least 4.2% of NYC cases due to multiple MPXV infections from distinct viral strains. Further, 13% of the sequences in infected patients had a lack of concordance between epidemiologic and genetic linkages. This study identified NYC-specific MPXV genomic profiles and provided a conservative estimate of simultaneous infections with multiple MPXV variants, likely due to behavioral risk factors during the peak of the outbreak in July 2022. Improving concordance between genomic and epidemiological clusters for mpox investigations may require expanding partnerships with testing labs, clinics, and community groups to enhance the representativeness of MPXV samples that are available for sequencing.

## Introduction

The causative agent of mpox in humans is the monkeypox virus (MPXV), a zoonotic *orthopox* virus [1]. MPXV causes symptoms that include fever, fatigue, headache, and a multi-stage rash that can spread between body sites. The genomic diversity of MPXV is represented in two clades, I and II. Clade I variants include sequences from Central Africa and are typically more transmissible and pathogenic. The first outbreak outside of the African continent was reported in the United States in 2003 [2] and belonged to Clade II, which includes two subclades, IIa and IIb and has reduced pathogenicity [3, 4]. Within Clade IIb, Lineages A and B emerged with subsequent sublineages [5]. In 2022, rapid transmission promoted the spread of MPXV which resulted in a global outbreak [4]. This outbreak was driven by MPXV lineage B.1, which gave rise to B.1.1 and other sublineages [5]. The first diagnosis of mpox in New York City (NYC) was on May 19, 2022 [6]. Since then, a total of 3,821 cases were reported in 2022 and more than 50 cases were reported in 2023 in NYC [7]. MPXV outbreaks have primarily spread through sex and other intimate contact among social networks of gay, bisexual, and other men who have sex with men (MSM) [8]. Risk factors for mpox-related hospitalization included immunosuppression and HIV co-infection [9].

Several genomic studies have been published on the 2022 outbreak in different countries around the world to understand the transmission and the evolution of the virus; however, most of these studies were limited by the number of genomes analyzed (n < 402) [4, 10–23]. Studying the intra-host genomic diversity of MPXV can provide additional resolution [4, 15] to study virus transmission networks. The intra-host genomic diversity of MPXV can be a result of multiple infections, viral evolution, or both within the host. Often, multiple infections in the same host can be identified on phylogenetic trees based on polyphyletic relations. However, multiple infections resulting from a close-knit transmission cluster involving related strains can be monophyletic on a phylogenetic tree [24]. Such relationships are usually over-represented for viruses like HIV among MSM communities and in people who inject drugs [24]. Noteworthy, 38-57% of mpox cases in the US are in people living with HIV [25] and may show similar patterns to HIV on a phylogenetic tree due to potentially similar behavioral risk factors.

MPXV has a slow mutation rate (1-2 substitutions per year) [26]; however, accelerated evolution (6 to 12-fold increase in mutation rate) has been documented for MPXV due to the genomic editing activities of host enzymes such as apolipoprotein B mRNA-editing catalytic polypeptide-like 3 (APOBEC3) [4, 17, 23, 27–29]. Newly acquired mutations can alter viral transmissibility [4] and reduce the efficacy of therapeutics and vaccines [30]. The probability of these mutations increases in cases of immunocompromised individuals, who may act as reservoirs for new variants [31].

An analysis of intra- and inter-host MPXV diversity is needed with a representative sample to enable a deeper understanding of MPXV evolution and transmission patterns. In this study, we used the largest genomic dataset to-date for MPXV infections in NYC (n=1,138 sequences from 758 total individuals). The sequencing data included 390 sequences collected from a single lesion from 390 individuals, and 748 sequences collected from two or more lesions from 368 individuals. Here we studied the genomic epidemiology of MPXV in NYC in comparison to MPXV genomes from around the world (n=2,968) using Nextclade and phylogenetic analysis. Mutational analysis was done to determine clade-defining mutations, APOBEC3 signatures, and the prevalence of clinically relevant mutations. Intra-host genomic diversity was assessed between different lesions within an individual, and inter-host genomic diversity was used to estimate co-infection rates in NYC. Finally, this study aimed to identify transmission networks for epidemiologically linked mpox cases in NYC using phylogenetics.

## Results

### Overview of MPXV genomes dataset

This study included 1,138 MPXV specimens collected between May 2022 and February 2023 and sequenced by the NYC Public Health Lab (PHL). The spatiotemporal distribution (**Supplementary Figures 1A and 1B**) shows a peak of specimen collection in July 2022 with the highest frequency of genomes sequenced from residents of Manhattan. Over half of individuals in this study (390/748) had single MPXV genomes collected from one lesion (34% of genomes) and the rest (358/748) had two or more MPXV genomes collected from two or more lesions (66% of genomes). Of the individuals with one genome, most of the sequenced specimens were from the genital area (62%, 241/390) (**Supplementary Figure 1C**). Similarly, for individuals with multiple genomes, the majority of sequenced specimens (54%, 404/748) were from the genital area, 10% (>68/748) were from upper body, and 5% (36/748) were from lower body sites. Specimens were collected from two lesions from the same anatomical site (*e.g.*, genital area) for 133 sequences and from different anatomical sites (*e.g.*, one lesion from the hand and one lesion from the genital area) for 468 sequences (**Supplementary Figure 1D**).

MPXV global genome sequences (n=2,968, excluding NYC sequences) included in this study were collected from 1985 through 2023 with representative sequences from five continents except Oceania. In this dataset, 2,878 genomes were from lineage B, 63 genomes were from lineage A, and 27 genomes represented the outgroup.

### Lineage assignment and phylogenetic analysis of MPXV genomes

In the combined NYC and global dataset, Nextclade designated 4,016 of the 4,106 sequences as lineage B.1 of MPXV clade IIb. In NYC, the most frequently observed B.1 sublineages were B.1.2, B.1.12, B.1.3 and B.1.7 (>50 sequences per sublineage) (**Figure 1A**). Nextclade lineage assignment had 98.9% and 98% agreement with the observed phylogenetic clades for sublineages observed in the global and NYC phylogenies, respectively (**Figure 1B**, **2A**). The phylogenetic clade that belonged to sublineage B.1.12 included predominantly sequences from NYC (∼93.6%) (**Figure 1B and Figure 2A**, red asterisk). For genomes designated as B.1 lineage by Nextclade, at least nine distinct clades were observed in the MPXV phylogeny with sequences in each clade predominantly from NYC and/or North America (**Figure 2B, Supplementary Figure 2, Table 1**).

**Figure 1.**
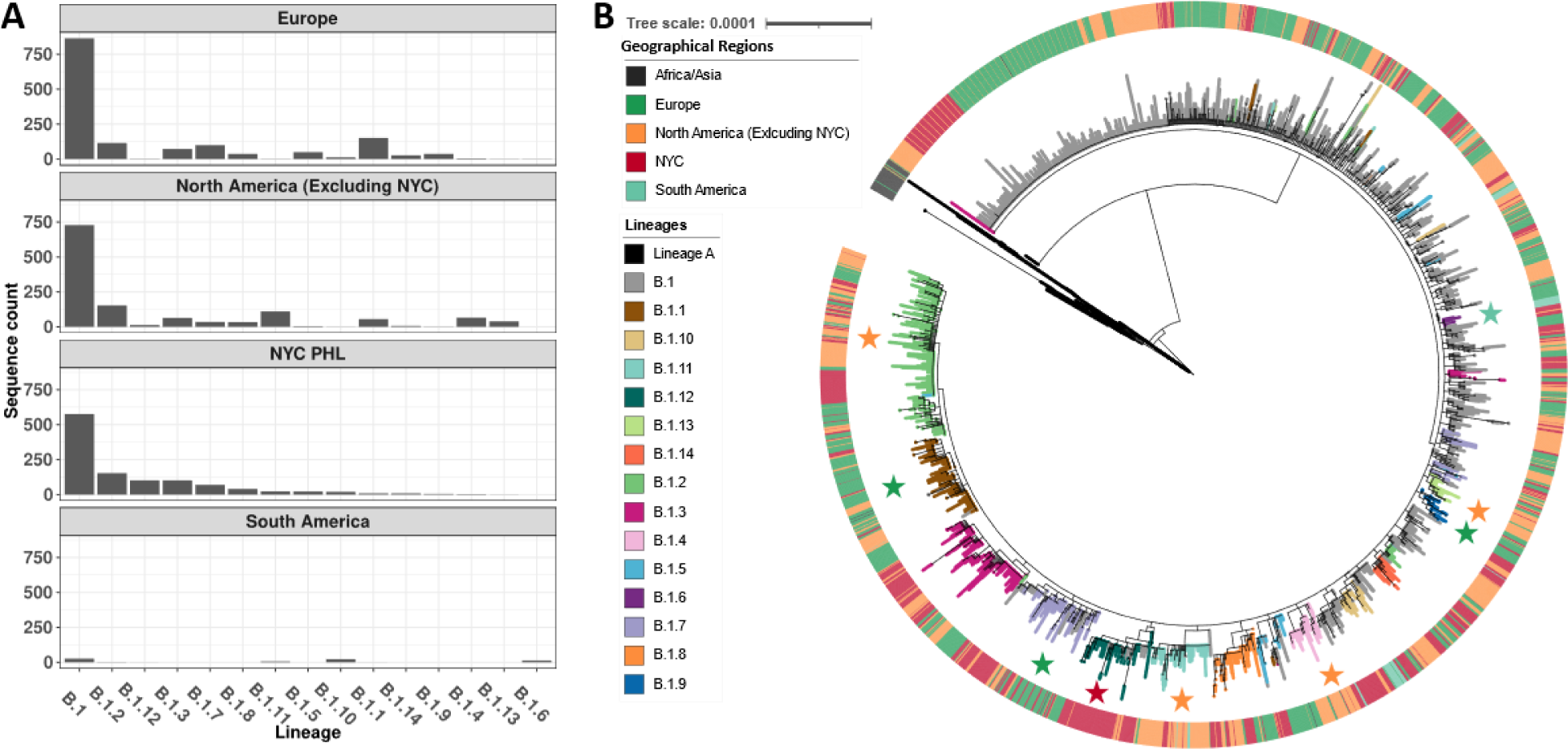
The spatial distribution and phylogenetic placement of MPXV lineages from global submissions. **A**) B.1 and sublineages assignment of the 2022 MPXV genome sequences from NYC, North America (Excluding NYC), Europe and South America. Sequences from the Africa and Asia were excluded from this figure due to small sample size (Africa: n=2; Asia: n=4). Lineage and sequence count per source were represented with x-axis and y-axis, respectively. Apart from the parent lineage B.1, the most frequently observed B.1 sublineages in Europe were B.1.1, B.1.2, B.1.7, B.1.3 and B.1.5 (>50 sequences per sublineage). In North America (excluding NYC), the most frequently observed B.1 sublineages were B.1.2, B.1.11, B.1.4, B.1.3, and B.1.1 (>50 sequences per sublineage). The most frequently observed NYC B.1 sublineages (>50 sequences per sublineages) were B.1.2, B.1.12, B.1.3 and B.1.7. **B**) Global phylogeny of MPXV genome sequences (including NYC sequences). The tree visualized with 4,078 sequences where 63 and 4,015 sequences were from lineage A and B, respectively. The collection dates ranged from 10/09/2017 to 01/09/2023. In the phylogeny, the branches were colored by Nextclade lineage assignment. The outer ring was colored by geographical region. Clades associated with specific geographical regions were shown using colored asterisks: Europe (green), North America (Excluding NYC, orange), NYC (red) and South Africa (cyan). All the B.1 sublineages were placed as distinct clades and had a 98.9% agreement with the Nextclade lineage assignment. Most sequences (93.6%) from sublineage B.1.12 were from NYC (red asterisk). Most sequences from sublineages B.1.11, B.1.13, B.1.2 and, B.1.4 were primarily from North America (excluding NYC) (orange asterisks); and sublineages B.1.1, B.1.7 and B.1.9 were primarily from Europe (green asterisks). B.1.6 sublineage was predominantly from South America (cyan asterisk).

**Figure 2.**
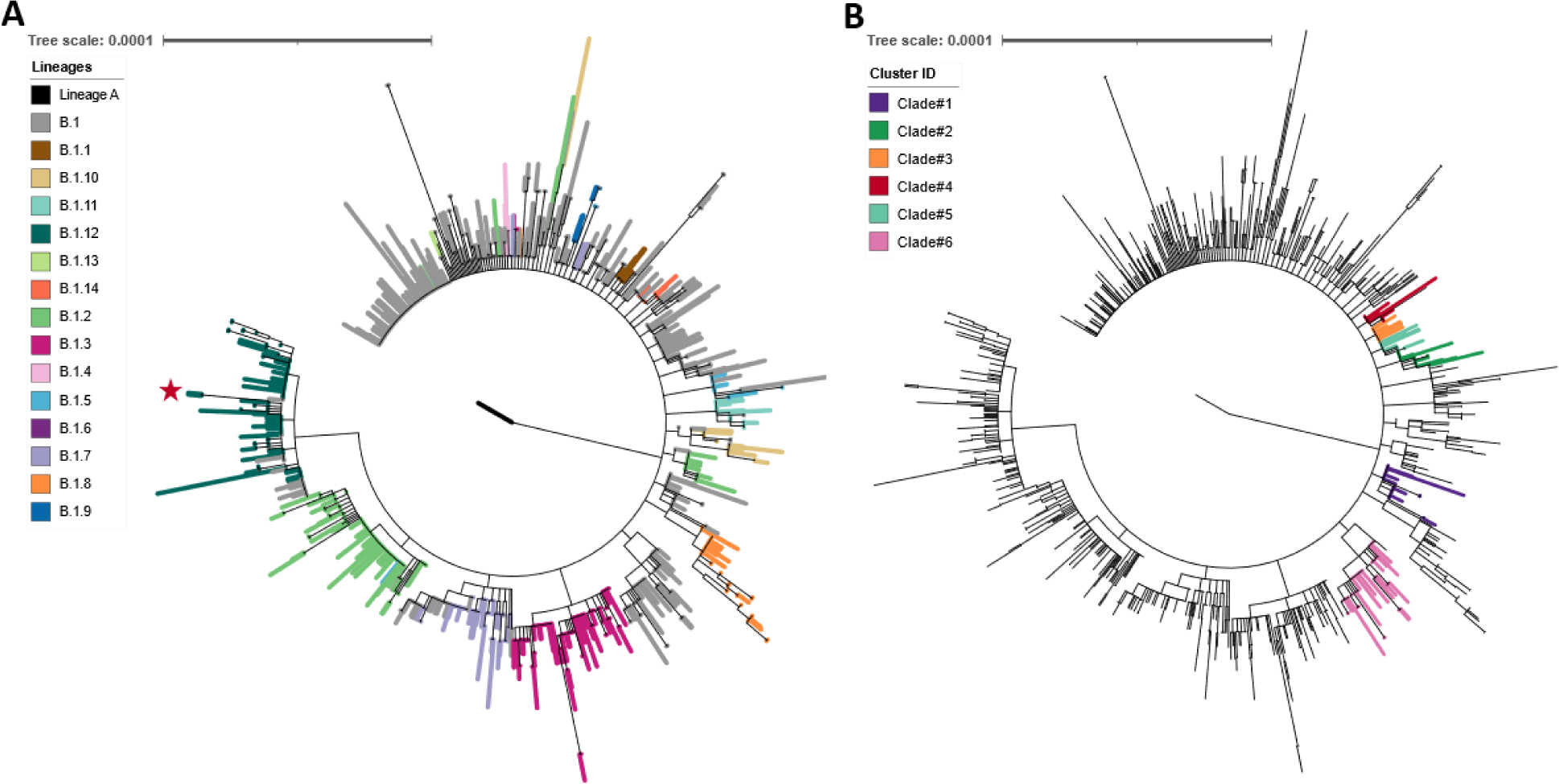
NYC MPXV genome phylogeny. **A**) A subtree of the global MPXV tree from Figure 1 containing 1,138 NYC sequences. The branches were colored by Nextclade lineage assignment. The B.1 sublineage clades were in 98.0% agreement with the Nextclade lineage assignment. The B.1.12 cluster was predominantly NYC specific (red asterisk). **B**) Clades emerged within B.1 assigned sequences. Lineage assignment was based on Nextclade. Six distinct clusters were observed in the NYC genome phylogeny (see **Supplementary Figure 2**) where sequences in each cluster were predominantly NYC specific and had at least one cluster-specific nonsynonymous mutation except for Cluster #6 (**Table 1**).

**Table 1.** Mutations in MPXV genomes with NYC and North America specific phylogenetic clades.

| ID | Position | Gene | Mutation (Amino acid change) | # Of Sequences | # Of NYC Sequences | # Of North American Sequences |
| --- | --- | --- | --- | --- | --- | --- |
| B.1.12 | 98233, 98455, 182950* | OPG117, OPG118, OPG210 | D729N, G4K, S532L | 119 | 102 | 13 |
| B.1.13 | 132520, 175093* | Intergenic, OPG204 | NA, D188N | 43 | 1 | 40 |
| B.1.4 | 34308* | OPG053 | S128L | 76 | 4 | 66 |
| Clade #1 | 36405 | OPG055 | A227T | 48 | 35 | 11 |
| Clade #2 | 39547, 152728 | OPG057, OPG180 | D217N, D13N | 23 | 22 | 1 |
| Clade#3 | 77462 | OPG098 | S188F | 53 | 17 | 36 |
| Clade #4 | 100867 | OPG119\|OPG120 | S161L | 17 | 15 | 2 |
| Clade #5 | 186177 | OPG210 | E1608K | 19 | 18 | 1 |
| Clade #6 | 43706, 124690, 141757 | OPG064, OPG145, OPG162 | Q731, S248, L36 | 68 | 59 | 9 |
| Clade #7 | 44960 | OPG064 | I313 | 48 | 1 | 47 |
| Clade #8 | 156429 | Intergenic | NA | 29 | 3 | 17 |
| Clade #9 | 103417, 150584, 173114 | OPG123, Intergenic, Intergenic | E552K, NA, NA | 28 | 0 | 28 |

\* Nextclade lineage defining mutation

### Lineage defining mutations, deletions, and the rate of MPXV evolution in NYC and North America

A total of 58 MPXV lineage-defining mutations were observed in lineage B, in which eight mutations were intergenic, 22 were synonymous, and 28 were nonsynonymous mutations. Most of these B lineage-specific mutations (∼91%) had APOBEC3 signatures (**Supplementary Table 1**). Genomic analyses showed that some B.1 sublineages harbored additional high frequency mutations that were not considered for defining lineage calls by Nextclade. For example, the B.1.13 clade observed in the MPXV global phylogeny had one high frequency intergenic mutation in position 132,520 (allele frequency=0.88) along with the lineage defining-mutation in position 175,093 (**Figure 1B**, **Supplementary Figure 3, Table 1**). Additionally, the NYC-specific lineage B.1.12 had two high frequencies nonsynonymous mutations in positions 98,233 (allele frequency=0.93) and 98,455 (allele frequency=0.93) along with the lineage-defining mutation in position 182950 (**Supplementary Figure 3, Table 1**). Genome sequences from six of the nine clades that were predominantly from NYC and/or North America had at least one clade-specific nonsynonymous mutation (**Table 1)**. The presence of these clade-specific mutations may qualify the sequences in those clades to be designated as new MPXV B.1 sublineages.

Deletions in surface glycoprotein *B21R* [12, 18] and the TNF receptor *crmB* [32] have been found in MPXV genomes sequenced from this outbreak; however, no deletions were found in these genes in the genomes sequenced by the NYC PHL. Other large-scale deletions were also rare in NYC sequences. Five occurrences of deletions were observed in 11 sequences collected from 10 individuals (**Table 2**). Sequences with a deletion from position 11,326:12,237/8 in the OPG023 gene were grouped into two distinct clades in the NYC phylogeny, likely due to two independent mutational events (**Supplementary Figure 4**).

**Table 2.** Deletions in NYC MPXV genome sequences.

| Deletion | Gene(s) | # of individuals |
| --- | --- | --- |
| 8197:9315 | OPG021 | 2 |
| 13543:16819 | OPG025-OPG027 | 1 |
| 11326:12237/8 | OPG023 | 6 |
| 149273:149608 | OPG174-175 | 1 |
| 159745:160509 | OPG185 | 1 |

The MPXV genome evolution rate was estimated to be 4.28e-5 subs/site/year using MPXV sequences collected from 2021 to 2023 (**Supplementary Figure 5**). Analysis of the amino acid changes in the phylogeny indicated that certain genes (OPG109, OPG110, OPG048) were more likely to have mutations (**Supplementary Figure 6**).

### APOBEC3 mutations in MPXV genomes

We assessed the prevalence of putative APOBEC3 signatures comparing the 2022 outbreak with previous years. The MPXV dataset was divided into three groups: (i) pre-2022 outbreak sequences retrieved from NCBI (1985-2021, n=81); (ii) 2022 global outbreak sequences, excluding NYC, retrieved from NCBI (n=2,887); (iii) 2022 NYC outbreak sequences sequenced by the NYC PHL (n=1,138). Compared to pre-outbreak sequences, both global and NYC MPXV sequences had significantly higher numbers of APOBEC3 signatures (*p*<0.001) (**Figure 3**). MPXV sequences in NYC also had slightly higher numbers of APOBEC signatures when compared to the global sequences during the 2022 outbreak (*p*<0.001). No significant differences were observed between any of these groups for non-APOBEC mutations.

**Figure 3.**
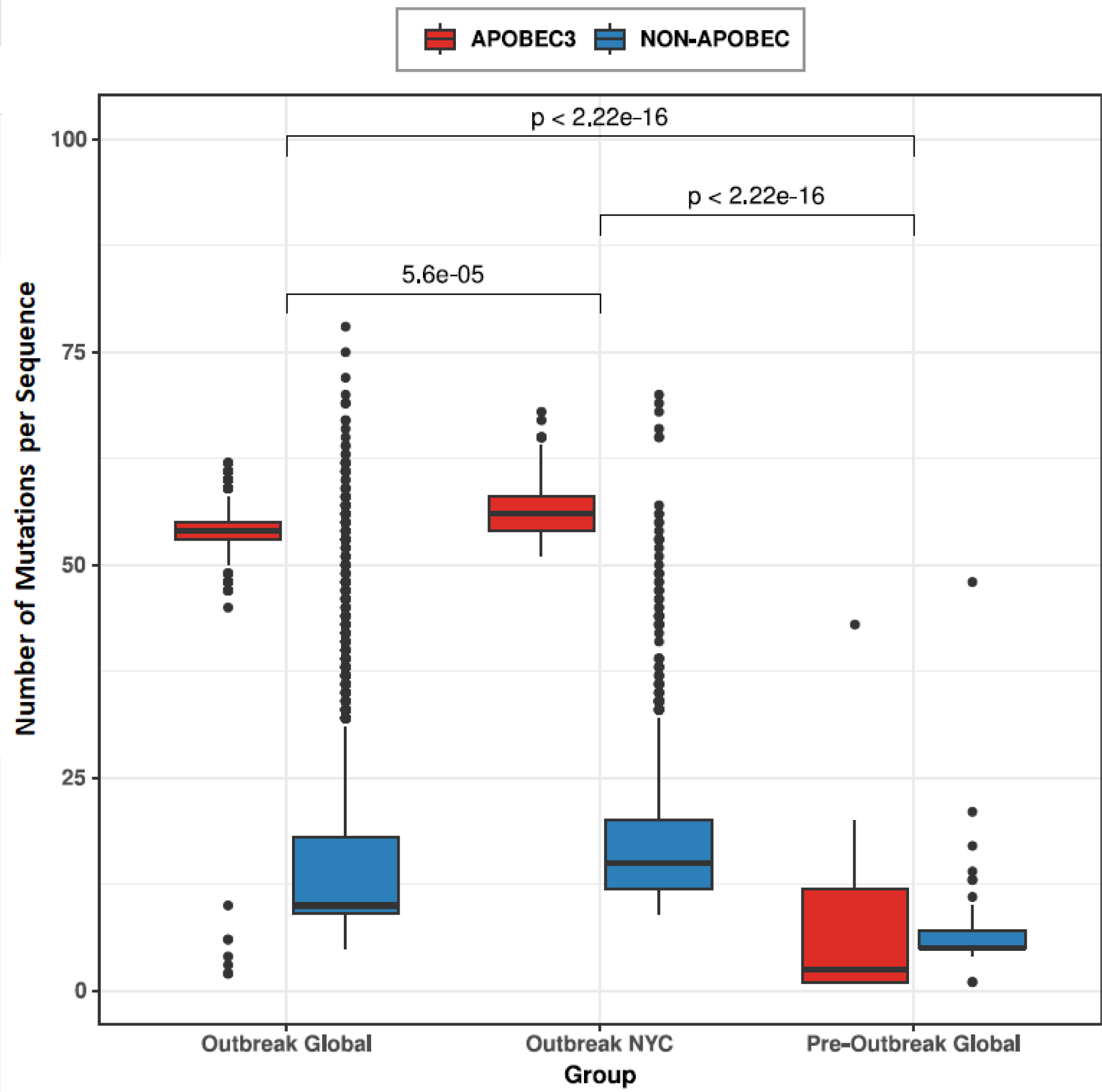
APOBEC3 signatures in MPXV genome sequences. The number of mutations with APOBEC3 signatures was significantly higher in the 2022 outbreak sequences (both global and NYC MPXV sequences) than the sequences from pre-outbreak period. The number of mutations per sample with and without APOBEC-3 signatures are shown on y-axis for: (i) Sequences retrieved from NCBI for MPXV genomes prior to the 2022 outbreak; (ii) 2022 outbreak global sequences excluding NYC; and (iii) 2022 NYC outbreak MPXV genomes sequenced by NYC PHL.

We screened 1,138 NYC MPXV genomes and found 446 (41.88%), out of the total of 1,065 mutations, had the “GA > AA” APOBEC3 signature. Four hundred twenty-five mutations (39.91%) had the “TC > TT” APOBEC3 signature, two (0.19%) had the “GG > AG” APOBEC3 signature, and 192 (18.03%) did not have any APOBEC signatures. Amongst the genomes screened, 51.5% of the APOBEC3 signatures occurred only once, 21.12% occurred twice and the remaining APOBEC3 signatures (27.38%) occurred more than twice.

### F13L mutations in NYC MPXV genome sequences

Mutations in the MPXV F13L gene homologue have been previously reported to be associated with tecovirimat (TPOXX) resistance. The NYC MPXV genomes showed a low frequency of F13L mutations (n=seven different mutations), four mutations of which were previously confirmed to be associated with TPOXX resistance [36] (**Table 3**). MPXV sequences with a TPOXX-associated mutation were from four individuals who had advanced HIV disease and were immunocompromised. These patients had been previously treated with TPOXX and were regarded as severe mpox cases. However, the risk of TPOXX resistance after treatment in the general population could not be evaluated due to a lack of information available on TPOXX treatment and outcomes in NYC.

**Table 3.** F13L mutations in NYC MPXV genomes.

| F13L mutations | TPOXX associated | # of individuals |
| --- | --- | --- |
| D217N | No | 14 |
| N267del | Yes | 1 |
| N267D | Yes | 1 |
| N267S | Yes | 1 |
| A290V | Yes | 1 |
| A288P | No | 1 |
| S369L | No | 1 |

### Infections with genetically distinct MPXV variants

When analyzing individuals for intra-host variation, we observed individuals that had multiple sequences with a high degree of variation between sequences (>10 SNPs). Amongst the individuals that had more than one lesion sequenced, 93.61% of individuals (337/360) had the same lineage assignment, 5.56% of individuals (20/360) had lineage assignments that were sublineages of the other sequence(s), and only 0.83% of individuals (3/360) had multiple genomes assigned as different lineages. To ensure the robustness of our multiple infection analysis, we chose only high-quality sequences in our NYC dataset and further masked genomic regions where many sequences had ambiguous bases. The final dataset used to infer a new phylogenetic tree specific to this analysis was 1,114 sequences.

Of the 360 individuals with MPXV genomes from multiple lesions, we identified 15 individuals whose genomes were polyphyletic in the phylogeny (*i.e.*, at least two distinct viral genomes mapped to disparate clades diverged at the root of the phylogeny; see Methods for details) (**Figure 4**). Eight additional sets of genomes were found to be distantly related but did not pass through the root (**Supplementary Figure 7**). The remaining 337 individuals had sequences that were monophyletic, direct ancestors (*i.e*., closely related, and consistent with potential intra-host variation), or closely related (*i.e.*, genomes did not have enough variation to be considered due to multiple infections). When considering only individual genomes where the branches separating the sequences pass through the root of the tree, we estimated 4.2% (15/360) of mpox cases had multiple infections with distinct MPXV variants in NYC. Expanding the definition to include the eight individuals with a node distance of at least four, the estimate for infection with multiple MPXV variants was 6.4% of mpox cases. These cases occurred in July 2022, the height of the outbreak in NYC (**Supplementary Figure 8**).

**Figure 4.**
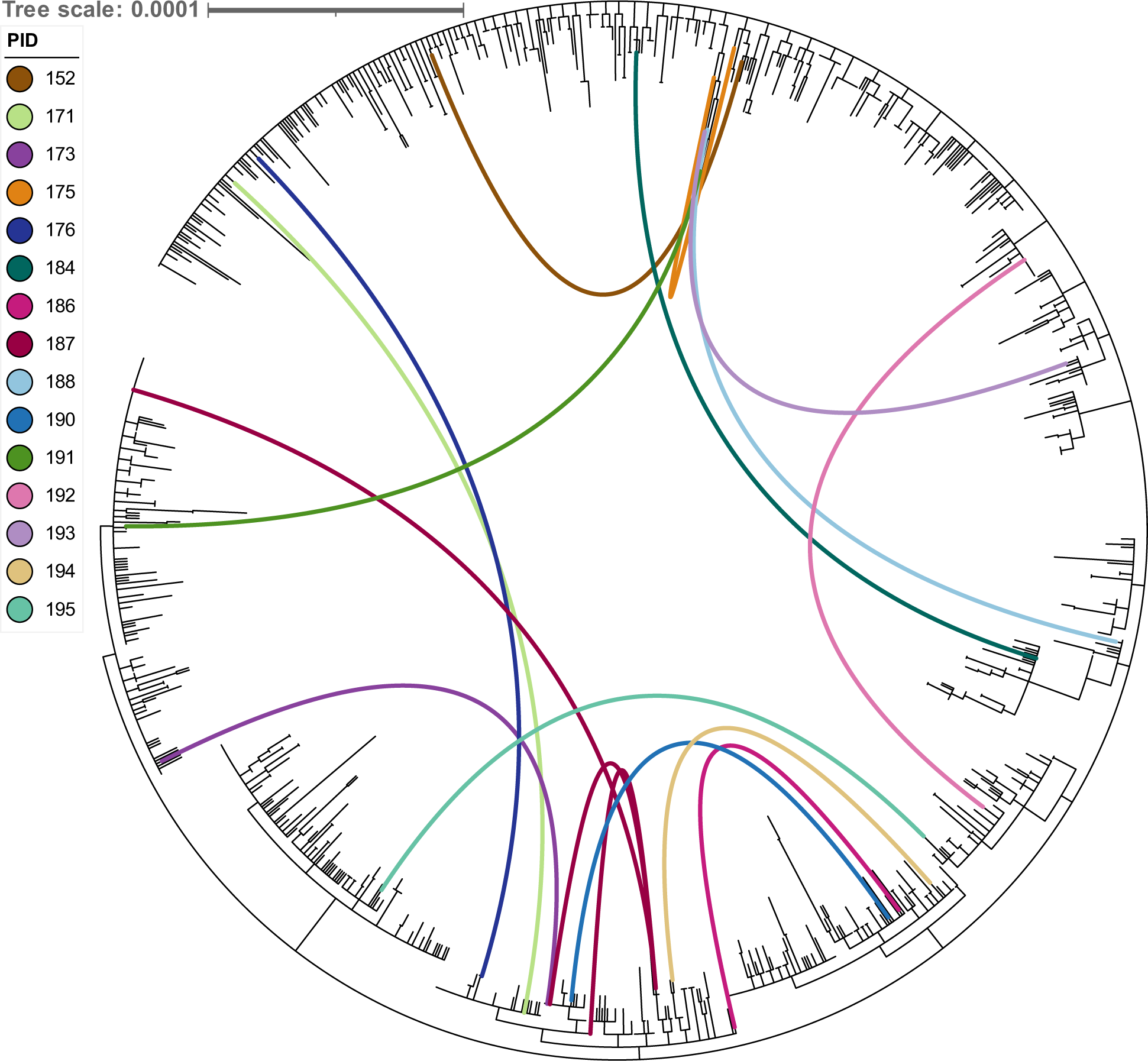
Infection with multiple MPXV variants in the 15 NYC individuals with sequences which diverged at root in the phylogeny. This NYC MPXV tree was inferred from 1,114 NYC sequences by masking the regions with low depth of coverage (**Supplementary Table 5**). Links in the figure were used to connect the sequences from the same individual that diverged at root using their phylogenetic placement. For Individual ID 187, one out of four sequences collected was found in a separate clade. The remaining three were found within the same clade, but one was distantly related by at least 4 nodes.

### Intra-host variation in NYC mpox outbreak specimens

The intra-host variation of NYC MPXV sequences was evaluated in 344 individuals with sequenced MPXV genomes from multiple specimens (*i.e.*, more than one lesion, majority sampled on the same day) and that were not due to multiple infections with distinct MPXV variants. We found that MPXV sequences from individuals with multiple lesions were identical in 172 individuals. Of the remaining 172 individuals, 119 individuals had multi-lesion sequences differing by only 1-2 mutations and 53 individuals had multi-lesion sequences differing by more than three mutations. Most intra-host variation had APOBEC3 signatures (**Figure 5A**). All the intra-host variation in 66.3% (114/172) of the individuals had APOBEC3 signatures, and at least 50% of the intra-host variation showed APOBEC3 signatures in 22.1% (38/172) of the individuals. Only 9.3% (16/172) of the individuals had intra-host variation that did not have APOBEC3 signatures (**Figure 5B**).

**Figure 5.**
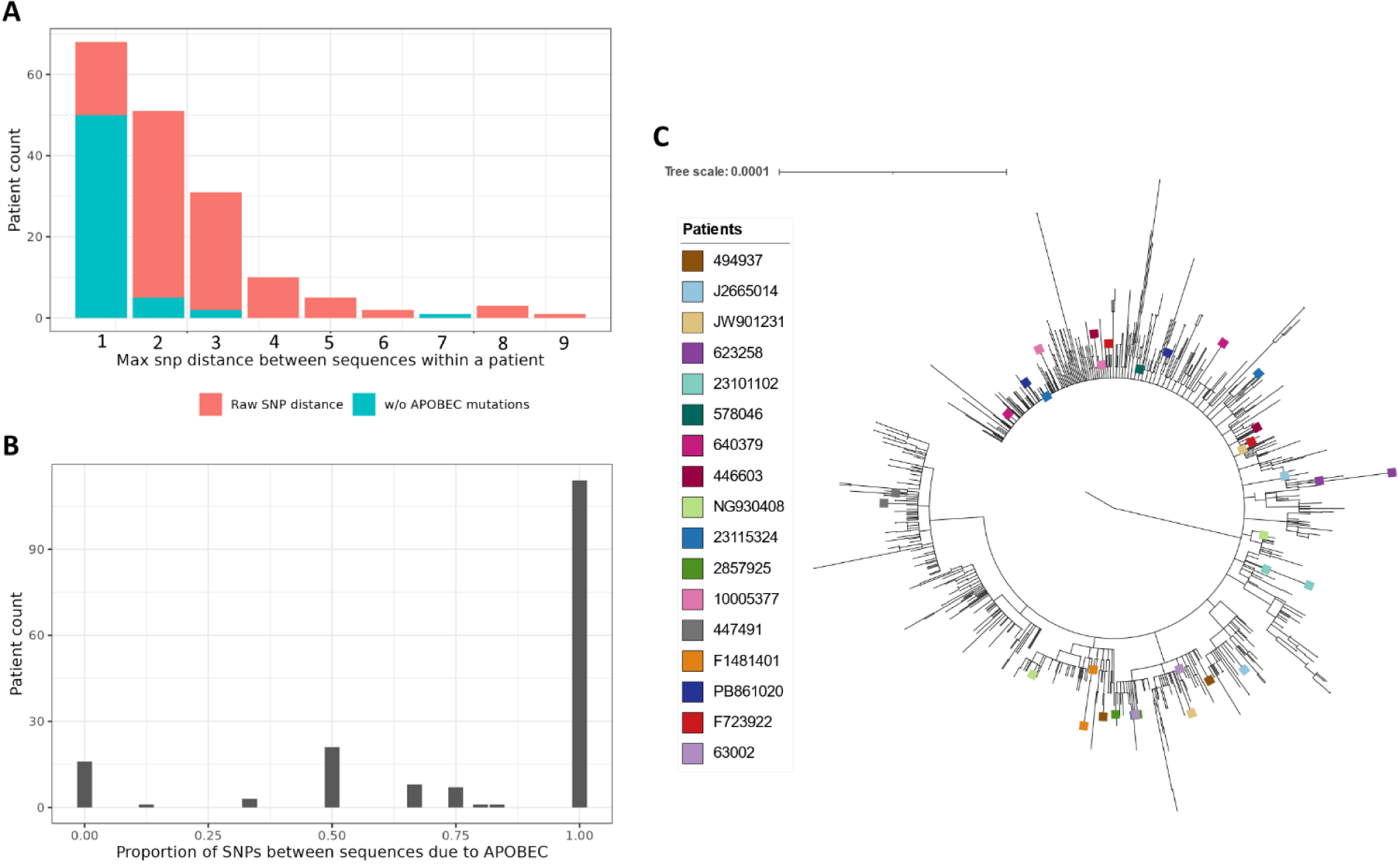
Intra-host MPXV genomic variation between sequences sampled from different lesions in the same individual. **A**) The frequency distribution of the maximum total SNP distance (with and without APOBEC3 signatures) between sequences within an individual. Out of 172 individuals, 119 (69.2%) individuals were closely related with 1-2 total SNPs between sequences, and 53 Individuals had sequences that were divergent with >=3 SNPs but a significant proportion of these mutational differences were due to APOBEC3. **B**) The proportion of intra-host SNP variation due to APOBEC. For 66.3% (114/172) of individuals, the observed mutational difference between sequences was completely due to APOBEC3. For 22.1% (38/172) of individuals, APOBEC3 contributed to 50% or more (but not 100%) of the mutational differences. Only 9.3% (16/172) of individuals had observed mutational differences that could not be attributed to APOBEC3. **C**) Intra-host phylogeny. Sequences from 17 individuals that had the greatest phylogenetic distance between them were annotated with colored tips and plotted on the NYC only phylogeny. Two individuals’ sequences (J2665014/light blue and NG930408/light green) resulted in non-monophyletic placements of sequences.

When these samples were evaluated on the NYC phylogeny, 21 individuals had samples with a genetic distance greater than 0.000021 substitutions/site. Sequences from the 17 individuals with the greatest within-host genetic distance are shown in **Figure 5C**, and the remaining four are shown in **Supplementary Figure 9**. Sequences from three individuals (two in **Figure 5C** and one in **Supplementary Figure 9**) were also found in different clades on the NYC phylogeny, illustrating that MPXV evolution within the same individual can be divergent enough to obscure the phylogenetic relationships between sequences. Although a rare occurrence (0.9% of individuals), this observation contrasts with Rueca et. al. [15] wherein longitudinally sampled individual sequences clustered together on a phylogenetic tree.

### Epidemiologically linked mpox cases in NYC

Contact tracing to identify epidemiologically linked mpox cases was performed for 43 individuals who were divided into 17 groups. Each group included epidemiologically linked pairs of individuals infected with mpox (**Table 4**, **Supplementary Table 2**). MPXV sequences from groups #1, #3, #5, #8 and #15 were genetically related (**Figure 6**, **Table 4**). Two MPXV sequences were genetically related in group #2 and five MPXV sequences were genetically related in group #6. The genetic relationship for the rest of the sequences from these two groups could not be resolved due to their placement in the basal part of the phylogenetic tree (**Figure 6**, **Table 4**). MPXV sequences from groups #7, #9, #11, #16, and #17 were not monophyletic but shared a common ancestor in the same clade and were therefore considered potentially genetically related (**Figure 6**, **Table 4**). MPXV sequences from groups #10, #12, #13 and #14 were placed in different clades and were not genetically related. Intra-host sequence pairs from groups #10, #12 and #13 were categorized as “Distantly Related” based on the multiple infection analysis performed in the previous section (**Figure 6**, **Table 4**). For example, group #12 included sequences from two lesions belonging to one individual. One of these intra-host sequences was assigned to the B.1 lineage (Accession #OQ469282) while the other sequence was assigned as B.1.7 (Accession #OQ469283). This observation suggests that sequencing only one lesion can result in inaccurate reconstruction of genomic transmission networks (**Table 4, Supplementary Table 2)**. Overall, 13.3% of sequences with epidemiological links were not genetically linked based on the phylogeny.

**Figure 6.**
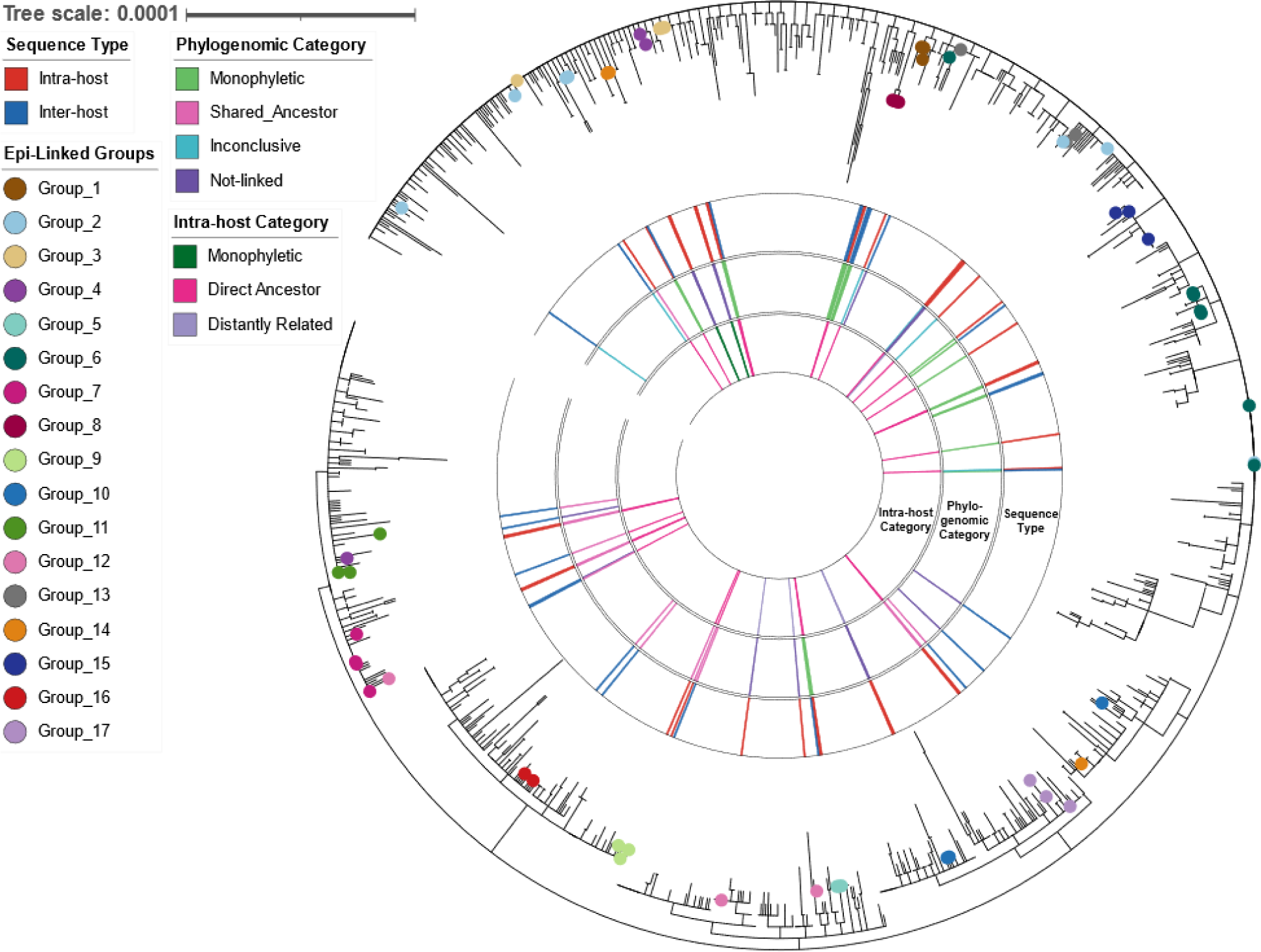
Epidemiologically linked MPXV cases in NYC. The symbols on the branches were colored by putative groups of epidemiologically linked cases reported by the Surveillance and Epidemiology Branch at the NYC Department of Health and Mental Hygiene. The outer ring (Sequence Type) in the figure was colored by the genome sequences that belonged to the same or different individual of epidemiologically linked cases in a particular group. The middle ring (Phylogenomic Category) is colored by the phylogenetic placement of the genome sequences of epidemiologically linked cases within group. Epidemiologically linked cases in groups with sequences placed on the same or sister branches were designated as “monophyletic”. Epidemiologically linked cases in groups for which genome sequences were in the same clade and shared a common ancestor with other NYC sequences in this clade were designated as “Shared Ancestor”. Epidemiologically linked cases in groups for which most of the genome sequences were placed in the basal part of the phylogenetic tree were designated as “inconclusive”. Epidemiologically linked cases in groups for which genome sequences were in different clades were designated as “Not-linked”. The inner ring (Intra-host category) was colored by the categories that were assigned using the phylogenetic placement of the genome sequences from the same individual (See multiple infection result section: Figure 4 and **Supplementary Figure 7**)

**Table 4:** Epidemiologically linked cases in NYC MPXV dataset.

| Epi-Linked Group | Accession #1 | Accession #2 | Sequence Type | Phylogenomic Category | Intra_host Category |
| --- | --- | --- | --- | --- | --- |
| Group_1 | OQ469130 | OQ468926 | Inter-host | Monophyletic |  |
| Group_1 | OQ504375 | OQ468926 | Inter-host | Monophyletic |  |
| Group_1 | OQ504375 | OQ469130 | Inter-host | Monophyletic |  |
| Group_2 | OQ469261 | OQ469262 | Intra-host | Inconclusive | Direct Ancestor |
| Group_2 | OQ469213 | OQ468568 | Intra-host | Inconclusive | Direct Ancestor |
| Group_2 | OQ468567 | OQ469261 | Inter-host | Inconclusive |  |
| Group_2 | OQ468567 | OQ469262 | Inter-host | Inconclusive |  |
| Group_2 | OQ468567 | OQ469036 | Inter-host | Inconclusive |  |
| Group_2 | OQ468567 | OQ469244 | Inter-host | Inconclusive |  |
| Group_2 | OQ468567 | OQ565521 | Inter-host | Inconclusive |  |
| Group_2 | OQ468568 | OQ469261 | Inter-host | Inconclusive |  |
| Group_2 | OQ468568 | OQ469262 | Inter-host | Inconclusive |  |
| Group_2 | OQ468568 | OQ469036 | Inter-host | Inconclusive |  |
| Group_2 | OQ468568 | OQ469244 | Inter-host | Inconclusive |  |
| Group_2 | OQ468568 | OQ565521 | Inter-host | Inconclusive |  |
| Group_2 | OQ469261 | OQ469036 | Inter-host | Inconclusive |  |
| Group_2 | OQ469261 | OQ469244 | Inter-host | Inconclusive |  |
| Group_2 | OQ469261 | OQ565521 | Inter-host | Inconclusive |  |
| Group_2 | OQ469262 | OQ469036 | Inter-host | Inconclusive |  |
| Group_2 | OQ469262 | OQ469244 | Inter-host | Inconclusive |  |
| Group_2 | OQ469036 | OQ469244 | Inter-host | Inconclusive |  |
| Group_2 | OQ469036 | OQ565521 | Inter-host | Inconclusive |  |
| Group_2 | OQ469244 | OQ565521 | Inter-host | Inconclusive |  |
| Group_2 | OQ469262 | OQ565521 | Inter-host | Monophyletic |  |
| Group_3 | OQ469140 | OQ469139 | Intra-host | Monophyletic | Direct Ancestor |
| Group_3 | OQ504392 | OQ469031 | Intra-host | Shared Ancestor | Direct Ancestor |
| Group_3 | OQ469031 | OQ469139 | Inter-host | Monophyletic |  |
| Group_3 | OQ469031 | OQ469140 | Inter-host | Monophyletic |  |
| Group_3 | OQ504392 | OQ469139 | Inter-host | Shared Ancestor |  |
| Group_3 | OQ504392 | OQ469140 | Inter-host | Shared Ancestor |  |
| Group_4 | OQ469032 | OQ469033 | Intra-host | Not-linked | Monophyletic |
| Group_4 | OQ469032 | OQ565498 | Inter-host | Not-linked |  |
| Group_4 | OQ565498 | OQ469033 | Inter-host | Not-linked |  |
| Group_5 | OQ469034 | OQ504394 | Intra-host | Monophyletic | Monophyletic |
| Group_5 | OQ469034 | OQ504393 | Inter-host | Monophyletic |  |
| Group_5 | OQ504393 | OQ504394 | Inter-host | Monophyletic |  |
| Group_6 | OQ468826 | OQ469163 | Intra-host | Inconclusive | Direct Ancestor |
| Group_6 | OQ419078 | OQ469231 | Intra-host | Monophyletic | Monophyletic |
| Group_6 | OQ468826 | OQ469160 | Inter-host | Inconclusive |  |
| Group_6 | OQ468826 | OQ419078 | Inter-host | Inconclusive |  |
| Group_6 | OQ468826 | OQ469231 | Inter-host | Inconclusive |  |
| Group_6 | OQ468826 | OQ469148 | Inter-host | Inconclusive |  |
| Group_6 | OQ468826 | OQ469232 | Inter-host | Inconclusive |  |
| Group_6 | OQ469163 | OQ419078 | Inter-host | Inconclusive |  |
| Group_6 | OQ469163 | OQ469231 | Inter-host | Inconclusive |  |
| Group_6 | OQ469163 | OQ469148 | Inter-host | Inconclusive |  |
| Group_6 | OQ469163 | OQ469232 | Inter-host | Inconclusive |  |
| Group_6 | OQ469160 | OQ419078 | Inter-host | Inconclusive |  |
| Group_6 | OQ469160 | OQ469231 | Inter-host | Inconclusive |  |
| Group_6 | OQ469160 | OQ469148 | Inter-host | Inconclusive |  |
| Group_6 | OQ469160 | OQ469232 | Inter-host | Inconclusive |  |
| Group_6 | OQ469163 | OQ469160 | Inter-host | Monophyletic |  |
| Group_6 | OQ419078 | OQ469148 | Inter-host | Monophyletic |  |
| Group_6 | OQ419078 | OQ469232 | Inter-host | Monophyletic |  |
| Group_6 | OQ469231 | OQ469148 | Inter-host | Monophyletic |  |
| Group_6 | OQ469231 | OQ469232 | Inter-host | Monophyletic |  |
| Group_6 | OQ469148 | OQ469232 | Inter-host | Monophyletic |  |
| Group_7 | OQ468605 | OQ468604 | Intra-host | Shared Ancestor | Direct Ancestor |
| Group_7 | OQ469159 | OQ504407 | Intra-host | Shared Ancestor | Monophyletic |
| Group_7 | OQ469159 | OQ468604 | Inter-host | Shared Ancestor |  |
| Group_7 | OQ504407 | OQ468605 | Inter-host | Shared Ancestor |  |
| Group_7 | OQ504407 | OQ468604 | Inter-host | Shared Ancestor |  |
| Group_7 | OQ469159 | OQ468605 | Inter-host | Shared Ancestor |  |
| Group_8 | OQ469047 | OQ468828 | Intra-host | Monophyletic | Monophyletic |
| Group_8 | OQ469047 | OQ469275 | Inter-host | Monophyletic |  |
| Group_8 | OQ468828 | OQ469275 | Inter-host | Monophyletic |  |
| Group_8 | OQ419004 | OQ469275 | Inter-host | Monophyletic |  |
| Group_8 | OQ469047 | OQ419004 | Inter-host | Monophyletic |  |
| Group_8 | OQ468828 | OQ419004 | Inter-host | Monophyletic |  |
| Group_9 | OQ468564 | OQ419075 | Intra-host | Shared Ancestor | Direct Ancestor |
| Group_9 | OQ468564 | OQ469268 | Inter-host | Shared Ancestor |  |
| Group_9 | OQ469268 | OQ419075 | Inter-host | Shared Ancestor |  |
| Group_10 | OQ565525 | OQ468537 | Intra-host | Not-linked | Distantly Related |
| Group_10 | OQ565525 | OQ468773 | Inter-host | Not-linked |  |
| Group_10 | OQ468773 | OQ468537 | Inter-host | Not-linked |  |
| Group_11 | OQ469293 | OQ468590 | Intra-host | Shared Ancestor | Direct Ancestor |
| Group_11 | OQ469293 | OQ469214 | Inter-host | Shared Ancestor |  |
| Group_11 | OQ469214 | OQ468590 | Inter-host | Shared Ancestor |  |
| Group_12 | OQ469283 | OQ469282 | Intra-host | Not-linked | Distantly Related |
| Group_12 | OQ469283 | OQ468587 | Inter-host | Not-linked |  |
| Group_12 | OQ468587 | OQ469282 | Inter-host | Not-linked |  |
| Group_13 | OQ468691 | OQ565515 | Intra-host | Not-linked | Distantly Related |
| Group_13 | OQ468691 | OQ565481 | Inter-host | Not-linked |  |
| Group_13 | OQ565481 | OQ565515 | Inter-host | Not-linked |  |
| Group_14 | OQ468873 | OQ468872 | Intra-host | Not-linked | Monophyletic |
| Group_14 | OQ468873 | OQ468606 | Inter-host | Not-linked |  |
| Group_14 | OQ468606 | OQ468872 | Inter-host | Not-linked |  |
| Group_15 | OQ468801 | OQ468662 | Intra-host | Monophyletic | Direct Ancestor |
| Group_15 | OQ468801 | OQ469191 | Inter-host | Monophyletic |  |
| Group_15 | OQ469191 | OQ468662 | Inter-host | Monophyletic |  |
| Group_16 | OQ565499 | OQ468660 | Inter-host | Shared Ancestor |  |
| Group_17 | OQ469213 | OQ468742 | Intra-host | Shared Ancestor | Direct Ancestor |
| Group_17 | OQ469213 | OQ419016 | Inter-host | Shared Ancestor |  |
| Group_17 | OQ419016 | OQ468742 | Inter-host | Shared Ancestor |  |

## Discussion

This study used the largest collection of MPXV sequences to date (1138 sequences from NYC and 2945 sequences for the global dataset) to understand the genomic epidemiology of MPXV in NYC. This study also included the largest dataset of sequences sampled from two or more lesions in the same individual (748 sequences). We found that some highly divergent sequences sampled from the same individual were non-monophyletic on the phylogenetic tree (**Figure 5C**). In addition, infection with multiple MPXV variants in the same individual occurred when case counts were high (**Supplementary Figure 8**). The estimation of infections with multiple MPXV variants in the same individuals in NYC was 4.2%, which is likely an under-estimation of the true prevalence. The phylogenetic analysis was based on consensus sequences derived from major alleles, and accounting for the minor alleles could likely have identified additional cases of infections with multiple MPXV variants. Incorporating within-host viral diversity in phylogenetic inference can improve the robustness of detecting clusters [37]. Therefore, future MPXV surveillance efforts can benefit from considering genomic variation outside of consensus sequences when attempting to identify potential transmission chains and clusters.

Four out of 17 groups with epidemiologically linked pairs of individuals infected with mpox had MPXV sequences that were not genetically related. The failure to link these individuals using phylogenetics analysis may be due to the inability to generate high quality MPXV genomes from available specimens or due to the lack of a representative sample of specimens that are available for sequencing. For example, individuals who had multiple partners might not have been tested at the NYC PHL and would not have been included in our analysis. Additionally, the epidemiologically linked cases did not distinguish between direct or indirect exposures (*e.g.*, attending common events). Nevertheless, it has been shown previously in HIV investigations among MSM in NYC that epidemiological linkage does not necessarily imply a genetic linkage consistent with viral transmission, and the percentage of named partners that had similar genomes varies depending on the number of partners and the perceived stigma of their behaviors [38]. Many of the groups with epidemiologically linked cases had unresolved genomic relationships, appearing highly related due to the slow evolving nature of MPXV genomes, which had minimal diversity during the early weeks of the outbreak. Thus, phylogenetics as a way to infer direct transmission needs to be approached with extreme caution because genomic similarity could be a result of slow evolution rather than transmission and genomic divergence could be a result of intra-host evolution mediated by APOBEC3 or infection with multiple MPXV variants within an epidemiological cluster.

The MPXV genome evolution rate was estimated to be 4.28e-5 substitutions/site/year using MPXV sequences collected from 2021 to 2023 (**Supplementary Figure 5**), which was consistent with a previous rate (5e-5 substitutions/site/year) estimated by analyzing 1,900 global MPXV genomes collected from 1958 to 2022 [33]. The MPXV genome evolutionary rate in this study was 4.8-fold higher than a previously reported rate (9e-6 substitutions/site/year) estimated using 87 MPXV genomes collected from 1978 until the early months of the 2022 outbreak. This discrepancy could be due to differences in collection date ranges and sample size. MPXV genomes had higher evolutionary rate in NYC comparison to MPXV genomes from around the world during the 2022 outbreak. Additionally, nine clades were identified with mutational profiles that were unique to NYC and North America (**Table 1**). Based on the phylogenetic placement and mutations, these clades were likely to be new MPXV sublineages that had emerged and spread within NYC or North America during the 2022 outbreak. Consistent with these findings, several unique lineages had emerged in the Midwest of the United States during the 2022 outbreak [19]. However, at the time of writing, these lineages were still classified as B.1 by Nextclade. The emergence of clades for subgroups of genomic sequences may present a challenge when using only Nextclade lineage assignment to determine genomic similarity or divergence.

Most of the genomic variations detected had APOBEC3 signatures (**Figure 3, Supplementary Table 1**), which is consistent with recent reports [4, 23, 28]. This APOBEC3-driven deamination has been observed with many DNA viruses and retroviruses and has been shown to be a driver for MPXV evolution during human-to-human transmission beginning in 2016 [35]. Despite this increased mutation rate, we found that the frequency of TPOXX-associated mutations [39] was low (7/1138) in our MPXV sequences. As a result, it is likely that additional factors, such as weakened immunity, contributed to these mutations in the four severe mpox cases. Therefore, information on the duration of infection, co-morbidities (*e.g.*, advanced HIV), or immunocompromised status can help better understand the virus’s genomic features [18]. There were several limitations in this study. On a global level, sequences from Africa and Asia were excluded from the global phylogeny due to the small sample size. On a local level, the NYC PHL was the sole testing facility for MPXV within the first seven weeks of the mpox outbreak in NYC. Specimen testing volume at PHL dropped once commercial lab testing was available. Despite these challenges, the number of individuals sequenced was reflective of the city’s epidemic curve [40]; and while randomly selected, sequences from global regions reflect the genomic diversity reported across the world [41]. Additionally, clinical data and time of infection (*e.g.*, acute, shedding, or recovery) was not available to study any associations between MPXV genomic characteristics and clinical outcomes. A potential bias in sample selection could be attributed to the lower real-time PCR Ct values required for sequencing (<30), which might have introduced bias towards a certain tissue, collection site, or phase of the infection (*e.g.*, macular, papular, vesicular, and pustular stages) where viral concentrations are higher. Most genomic sequences were collected from the genital region. As a result, it is unclear how well genomic variations of MPXV can be captured in other specimen types where viral concentration is lower (*e.g.*, blood, saliva, oral/rectal swabs) [42]. Longitudinal sampling from the same lesions and hosts could have also provided valuable information on the intra-host evolution of the virus. However, intra-host variations that were observed in this study were primarily from samples from the same individual collected during the same day. The multiple infection analysis was therefore focused on the genomic diversity within an individual at the time of diagnosis. Last, but not least, epidemiological contact tracing data maybe incomplete due to perceived stigma related to the number of named or unnamed partners and sexual behaviors [38].

In conclusion, this study included the largest genomic dataset collected during the 2022 mpox outbreak, featuring the biggest archive of specimens from the same individuals. The integration of genomic and epidemiological data revealed transmission relationships involving individuals infected with multiple MPXV variants. Analyzing infections with multiple MPXV variants showed that individual lesions often do not represent the complete diversity of within-host variation and may have distinct genomic profiles for MPXV. Some of the exposures from contacts documented through traditional epidemiological methods were not supported by the viral genomic diversity between individuals, which may have been underrepresented due to a variety of biological as well as technical and sampling reasons. Improving concordance between genomic and epidemiological clusters may require improving the reconstruction of high-quality genomes from different lesion sites as well as improving the representativeness of collected samples that are available for sequencing by expanding partnerships with testing sites and collaborations with community groups, which may be a challenge during resource-constrained times. Results from this study highlight the importance of gathering thorough epidemiological, clinical, and case data for outbreak investigations followed by thorough and careful interpretation of the data to perform genomic epidemiology for MPXV.

## Methods

### Specimen selection

Lesion swabs from one or more body sites from suspected mpox cases were submitted to the New York City Public Health Laboratory (NYC PHL) for testing using Non-Variola *Orthopoxvirus* (NVO) Real-Time PCR. DNA was extracted from lesion swabs or swabs in viral transport media using manual or automated DNA extraction platforms and the QIAGEN QIAamp® DSP DNA Blood Mini Kit or the QIAGEN EZ1&2 DNA Tissue Kit, respectively. NVO detection was performed using the CDC Laboratory Response Network (LRN) Non-Variola *Orthopoxvirus* real-time PCR Assay targeting the E9L gene target found in MPXV. Specimens with a Ct value < 37 were considered positive for Non-Variola *Orthopoxvirus* DNA. Specimen with Ct values <= 30 were sequenced after DNA extraction.

### Whole genome sequencing

NYC PHL utilized PrimalSeq [43] for MPXV whole genome sequencing. PrimalSeq for MPXV is a tiling PCR approach which consisted of 163 MPXV-specific primer pairs divided into two primer pools for the initial PCR step. These primers spanned nearly the entire length of the genome (from base 356 to 196424) and produced overlapping amplicons with an average length of approximately 2000 bp. These amplicon pools were cleaned of primers using Beckman Coulter Ampure XP beads (Cat. # A63880). The bead-cleaned amplicons were subjected to a tagmentation-based library preparation for small PCR amplicon input using the Illumina DNA Prep Kit (Cat. # 20018705) with IDT for Illumina Unique Dual Index Sets (Cat. #s: 20027213(UD-A), 20027214(UD-B), 20042666(UD-C), 20042667(UD-D)). The libraries were run on an Illumina MiSeq at 151bp paired-end reads with a llumina MiSeq Reagent Kit 600-cycle v3 kit (Cat. # MS-102-3003) or a NextSeq2000 with a NextSeq 1000/2000 P2 Reagent 300-cycle kit v3 kit (Cat. # 20046813).

### Assembly and variant calling of NYC PHL MPXV outbreak genomes

Nextera DNA Flex CD Indexes were removed as part of FASTQ generation. Reads were quality trimmed using Trimmomatic 0.36 [44] before mapping to the MPXV reference genome NC_063383.1 using minimap2 v2.17-r941 [45]. Samtools v1.13 [46] was used to sort and index mapped reads before trimming primer sequences from alignment files with iVar [47]. Variant Call Files (VCF) were created by BCFtools using a minimum quality score of 20, depth of coverage (DP) of 20, and frequency threshold of 0.65 from the primer trimmed alignment files. The primer trimmed alignment files were also used to generate consensus sequences with iVar consensus with the same minimum quality score, DP, and frequency thresholds that were used to generate the consensus sequences. For phylogenetic analyses, **S**ome **Qui**ck **R**earranging to **R**esolve **E**volutionary **L**inks (squirrel) (https://github.com/aineniamh/squirrel) was used to mask the inverted terminal repeat (ITR) regions as well other problematic regions. Sequences with a genome coverage <90% were excluded from downstream analyses, resulting in a total of 1,138 MPXV genomes from NYC PHL.

### Curation of global MPXV genome sequences dataset

Global MPXV genome sequences (n=2,968, excluding any sequences from NYC) and associated metadata were obtained from the National Center for Biotechnology Information (NCBI) virus database (https://www.ncbi.nlm.nih.gov/labs/virus/vssi/#/) on February 1^st^, 2023. Partial sequences and/or sequences with missing collection dates and geographic origins were excluded from downstream analyses. The complete list of 1,138 NYC PHL and 2,968 global MPXV sequences is available in **Supplementary Tables 3 and 4**.

### Lineage assignment

The lineages of global and NYC MPXV genome sequences were assigned using Nextclade (https://clades.nextstrain.org/) with MPXV reference genome NC_063383.1 (collected from Nigeria, August 2018) and Dataset name hMPXV. The combined dataset which contains 4,106 MPXV sequences was aligned to the reference using MAFFT v7.490 [48] with the --6merpair option. A maximum-likelihood tree was inferred by IQTree2 [49] with the GTR+F+I model of nucleotide evolution and NNI search option. Trees were visualized using iTOL (https://itol.embl.de/) [50]. MPXV genome divergence rate was calculated using Nextstrain [51] with the reference genome and the previously generated IQTree2 maximum-likelihood tree. A time-resolved tree was obtained, but many samples were filtered out by the --clock-filter-iqd 4 option. This led to only sequences from 2021 onward to be included for a total of 3,936 sequences.

### Characterization of MPXV genomic variants, deletions and TPOXX resistance

Genome variants in MPXV sequences were called using NucDiff [52] against the reference genome, and the resulting VCFs were annotated with SnpEff [53]. Ambiguous bases (N) were excluded from further analyses. Variant analyses were performed using a custom-written R script and visualized in R with ggplot2. Potential APOBEC3 signatures in the MPXV genomes were detected using Mutation profile (https://github.com/insapathogenomics/mutation_profile) [4]. To identify clade-specific mutations and other clinically relevant mutations, we restricted our analysis to genomes sequenced at the NYC PHL due to limitations in the availability of alignment and VCF data from the global submissions. To detect deletions, Samtools depth was used to flag regions of the genome with a DP of 0. To reduce false positives, deletion regions were determined to be amplicon dropouts and discarded if the surrounding regions had a DP less than 20. Furthermore, the remaining deletion regions were manually reviewed using IGV [54]. TPOXX (tecovirimat) resistance associated mutations were identified using SnpEff and searching for F13L mutations listed in the FDA Microbiology Review on TPOXX [39].

### Intra-host variations in NYC mpox outbreak specimens

snp-dists (https://github.com/tseemann/snp-dists) was used to determine the nucleotide differences between sequences from the same individual (*i.e.*, differences in ambiguous bases are ignored). Sequences that were divergent enough to be considered potential multiple infections were excluded from this analysis. Genomic variation between sequences from the same individual were then annotated as having an APOBEC signature or not. Phylogenetic distance between these sequences was determined using the R package adephylo and manually assessed by plotting on the NYC specific phylogenetic tree.

### Inferring Infection with multiple MPXV variants

#### Selecting sequences to identify multiple infections and genome masking

Sequences with more than 5% N in the squirrel masked [55] sequences were excluded, resulting in 1,114 sequences from 742 individuals. Most individuals (n=382) had only a single individual sequenced MPXV genome; however, 360 individuals had MPXV genomes sequenced from at least 2 lesions (n=732 genomes).

To minimize sequencing artefacts related to batch effects, amplicon dropout, sequencing error, or the 3’ terminal repeat region [56], the proportion of sequences with genome positions with low DP (<20) at each position was calculated to identify potential regions to mask. Using k-means in R with three centers starting with 25 random centers, positions were categorized into three groups of proportions: low (0.1 – 9.16%), medium (9.25 – 51.9%), high (52.1 – 100%). Positions classified as medium or high were used to identify larger regions to mask.

A BED file for masking the 1,114 genomes was created by including all regions of medium or high proportions of low DP except the k-means group of regions with the smallest stretch of low DP. Additionally, the 3’ inverted terminal repeat (ITR) was masked due to poor read mapping quality (MAPQ). Genomic regions that were masked due to potential low-quality sequences and the 3’ ITR are listed in the **Supplemental Table 5**. In total, 12 regions were masked, resulting in high quality genome coverage between 89.91% to 93.67% in the NYC dataset for the multiple infection analysis.

#### Phylogenetics

After masking, a phylogenetic tree comprising only MPXV genomes sequenced by PHL (n=1114 genomes) was inferred using IQ-Tree2 version 2.1.3 under a GTR+F+I substitution model, with a minimum branch length of 1e-10 substitutions/site and collapsing polytomies. For the 360 people with more than one sequenced viral genome, we determined the phylogenetic relationship and distance among these viruses using the ete3 python package [57]. Genetic distance was assessed using HIV-TRACE [58]. Individuals were potential multiple infection with different MPXV strains if at least two of their sequenced viruses were (i) separated by at least 3 nucleotide substitutions, (ii) not each other’s direct ancestor or descendant, (iii) were separated by at least four internal nodes in the phylogeny, (iv) were at least one nucleotide substitution from the basal polytomy at the root, and (v) shared a most recent common ancestor (MRCA) at the root of the phylogeny. As a sensitivity analysis, we also considered multiple infection if all these criteria were met except for sharing an MRCA at the root of the phylogeny.

### NYC epidemiologically linked mpox outbreak specimens

Whole genome sequencing, lineage assignment and phylogenetic analyses were performed to confirm epidemiologically linked mpox outbreak cases in which the individuals had contact with one or more persons who had mpox and transmission by the usual modes of transmission was deemed likely. Lineage assignment, phylogenetic inference and visualization were performed using NextClade, IQTree2 and iTOL (https://itol.embl.de/), respectively.

## Supporting information

Supplementary_Table_2_Epilinked_results

Supplementary_Table_3_&_4_MPXV_genome_sequences

## Acknowledgements

The authors acknowledge the Grubaugh Lab at Yale University for their help in providing the primers for the initial proof of concept.

This publication is supported by Cooperative Agreement Number NU60OE000104 (CFDA #93.322), funded by the *Centers for Disease Control and Prevention (CDC) of the US Department of Health and Human Services (HHS)*. Its contents are solely the responsibility of the authors and do not necessarily represent the official views of APHL, CDC, HHS or the US Government. This project was 100% funded with federal funds from a federal program of $120,402,978.

## Data availability statement

All data used in this study are publicly available. NYC MPXV sequence data are available on NCBI (Bioproject: PRJNA949682, **Supplementary Table 3**). Sequencing data that were retrieved from NCBI for the Global Phylogeny are summarized in **Supplementary Table 4**.

