## Supplementary material for "Genomic Epidemiology of Monkeypox Virus During the 2022 Outbreak in New York City": Supp_Figures_and_Tables

### Supplementary Figures

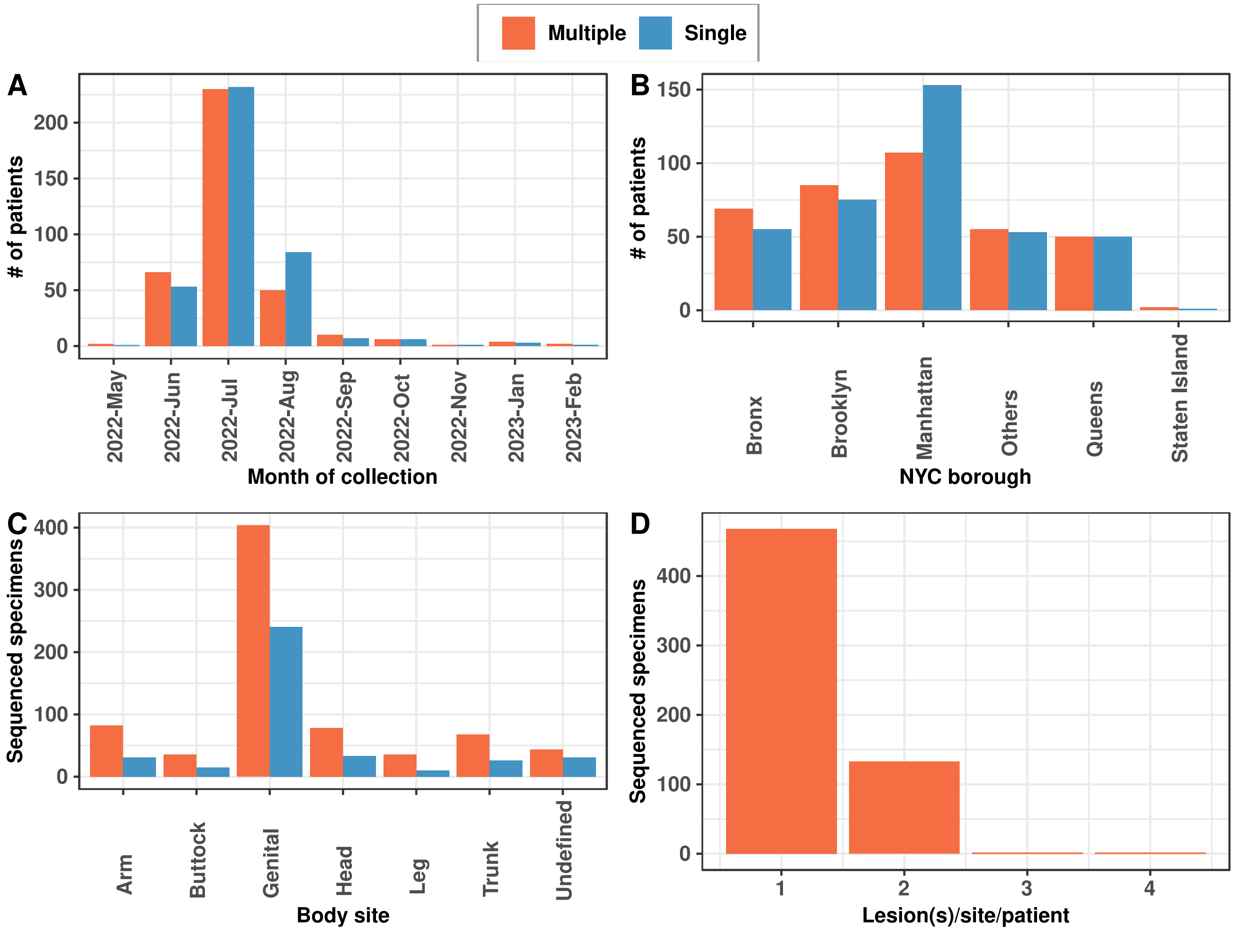


**Supplementary Figure 1. Characteristics of MPXV genomes sequenced by the NYC PHL during the 2022 outbreak. A**) Temporal distribution of the number of individuals from whom specimens were collected and genomes were sequenced. **B**) Distribution of the number of individuals by NYC borough of residence from whom specimens were collected and sequenced. **C**) Tissue distribution of sequenced genomes from specimens collected. **D**) Summary of the number of sequenced specimens per individual.

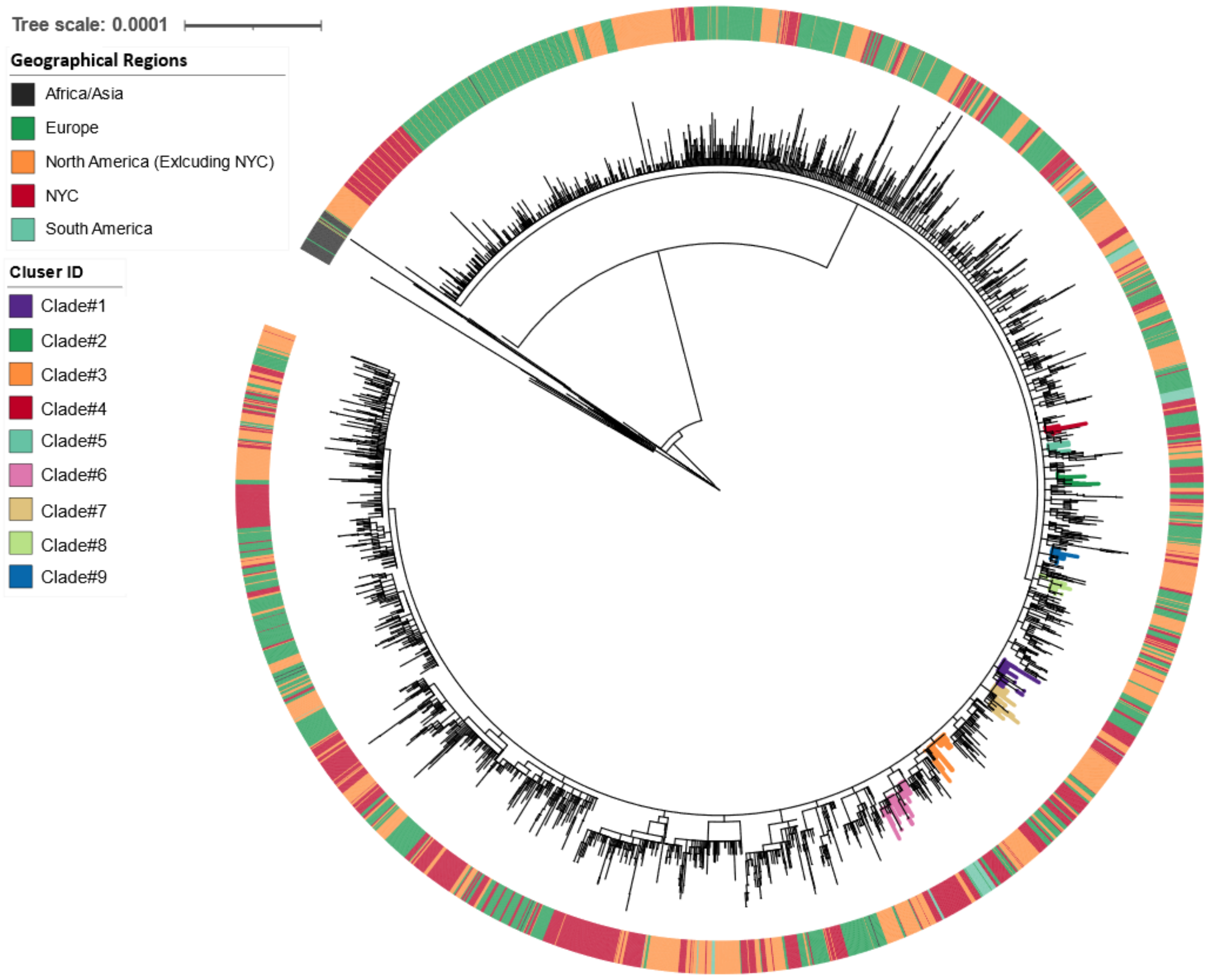


**Supplementary Figure 2**. **Clades in the MPXV genome global phylogeny**. For sequences with B.1 Nextclade lineage assignment, nine distinct clades were observed in the MPXV genome phylogeny. The branches were colored by Clade ID. The outer ring in the figure was colored by geography. Sequences in each clade were predominantly NYC and/or North America specific and had at least one clade-specific mutation (**Table 1**).

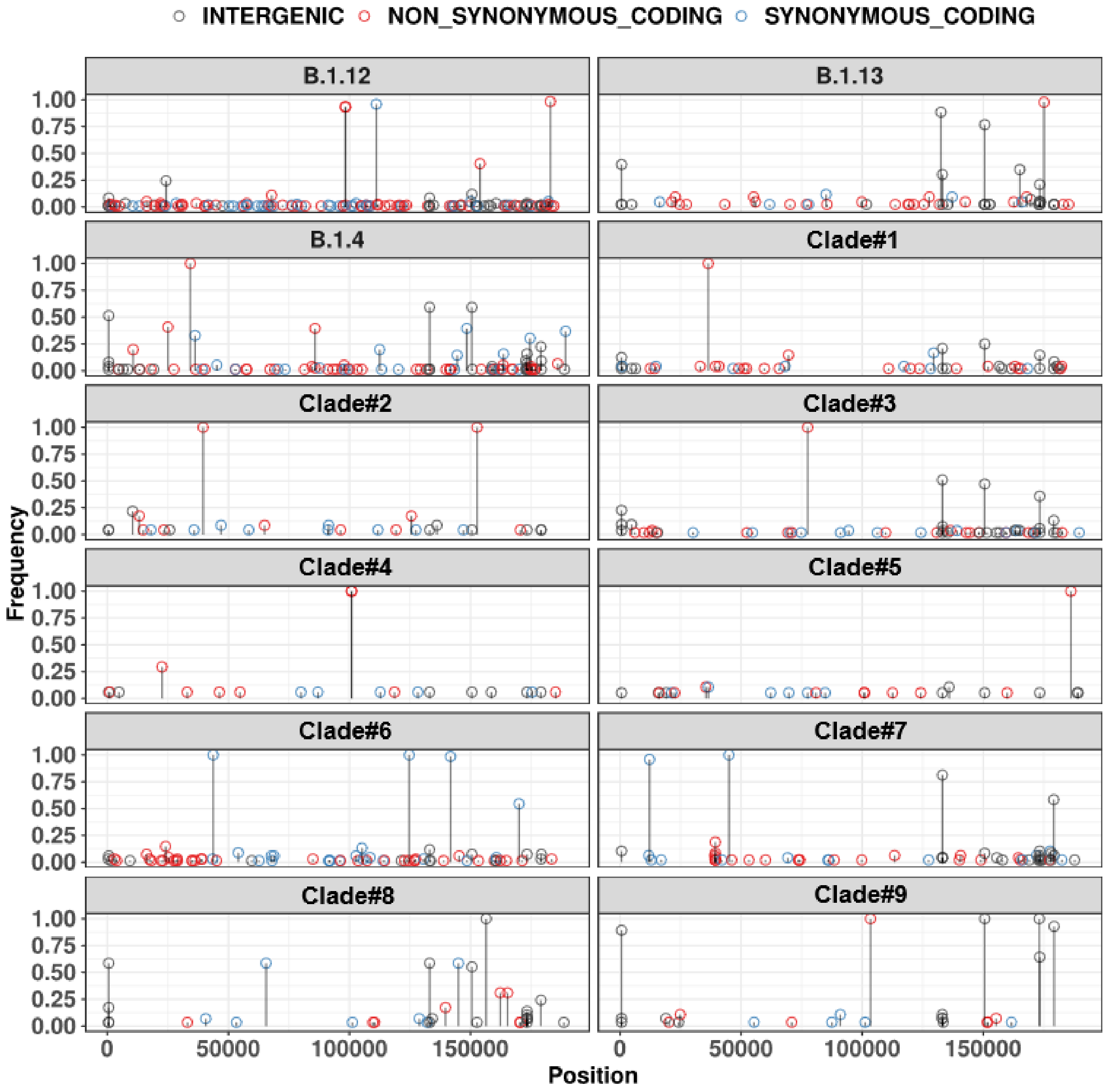


**Supplementary Figure 3**. **Mutations in B.1 sublineages and newly observed phylogenetic clades that were specific to NYC and North America**. Genome position and proportion of the mutations were represented by x-axis and y-axis, respectively. Each sublineage and clade had its unique lineage defining mutations (See **Table 1**). NYC specific sublineage B.1.12 had two additional high frequency nonsynonymous mutations in position 98233 and 98455 (left panel, row 1). Nine distinct clades with predominantly NYC and/or North America genome sequences were observed in the MPXV global and NYC genome phylogeny (**Figure 1B**, **Supplementary Figure 2**). Clade #1, Clade #3, Clade #4, Clade #5, and Clade #9 had one nonsynonymous clade defining mutation in position 36405, 77462, 100867 186177 and 103417 respectively. Clade #2 had two nonsynonymous clade defining mutations in position 39547 and 152728 (See **Table 1**).

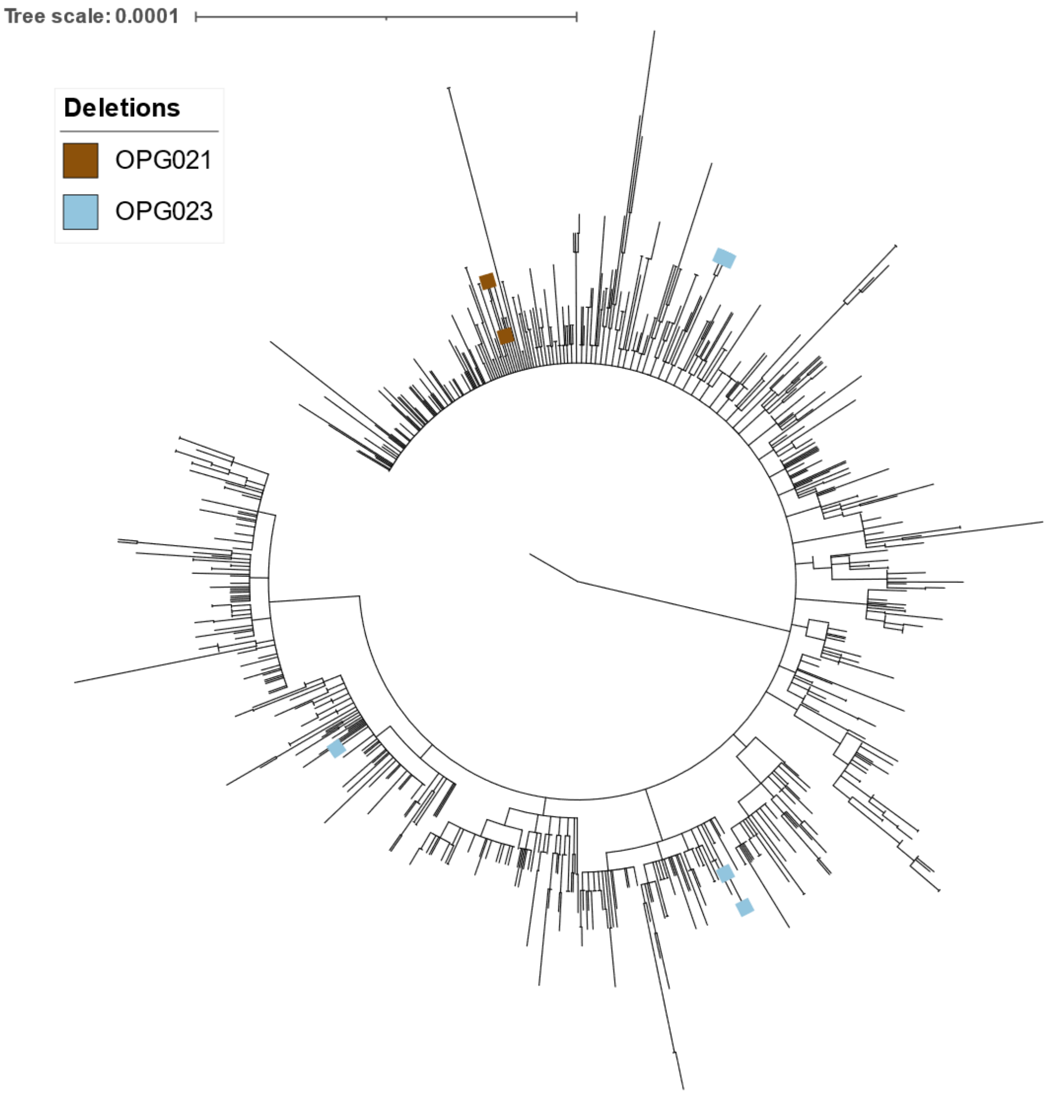


**Supplementary Figure 4**. **Phylogenetic placement of sequences with deletions in two positions.** Deletions shown are positions 8197-9315 (brown) in gene OPG021 and 11326-12237/8 (blue) in gene OPG023. The sequences with deletion 11326-12237/8 were grouped into two distinct clades in the NYC only phylogeny.

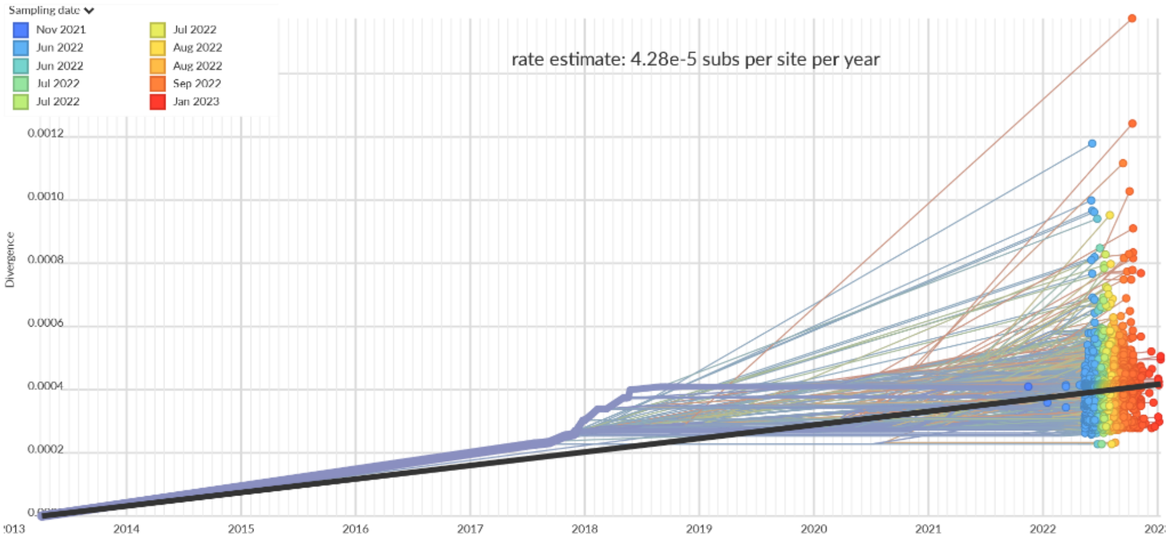


**Supplementary Figure 5**. **MPXV rate of divergence**. MPXV genome evolution rate was estimated to be 4.28e-5 substitutions/site/year by performing regression analysis using genome divergence from reference (y-axis) versus collection date (x-axis).

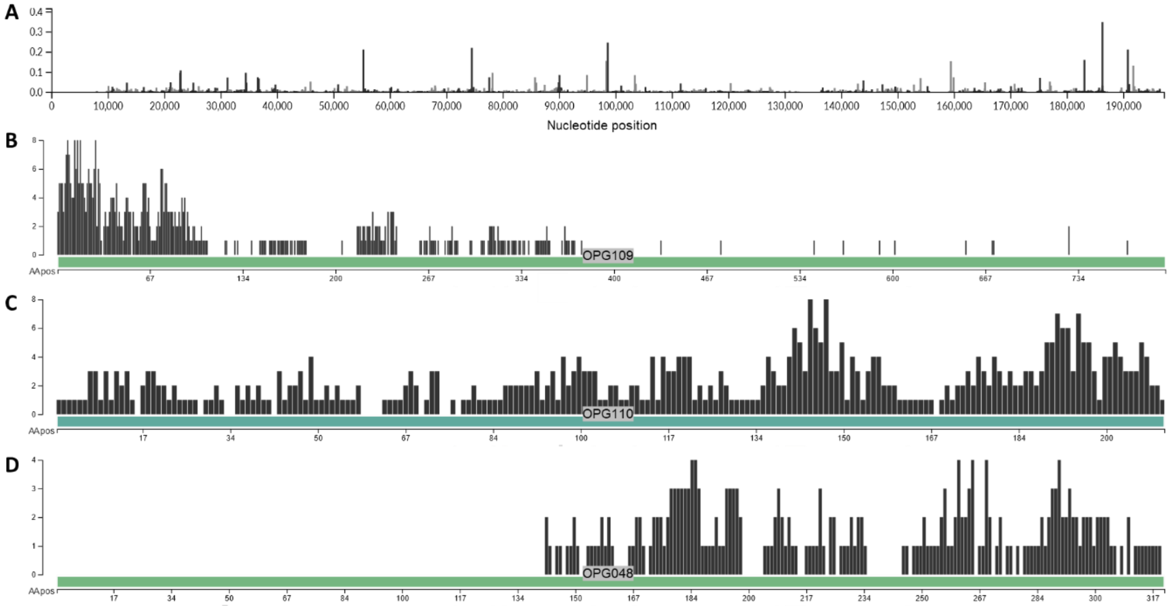


**Supplementary Figure 6**. **Event plot of amino acid changes in MPXV Nextstrain build**. **A**) Ancestral state reconstruction was used to infer the number of changes that occurred on the phylogeny (y-axis) at each nucleotide position (x-axis). Genes with the highest number of mutational events were **B**) OPG109 (532 total events), **C**) OPG110 (463 total events), and **D**) OPG048 (235 total events).

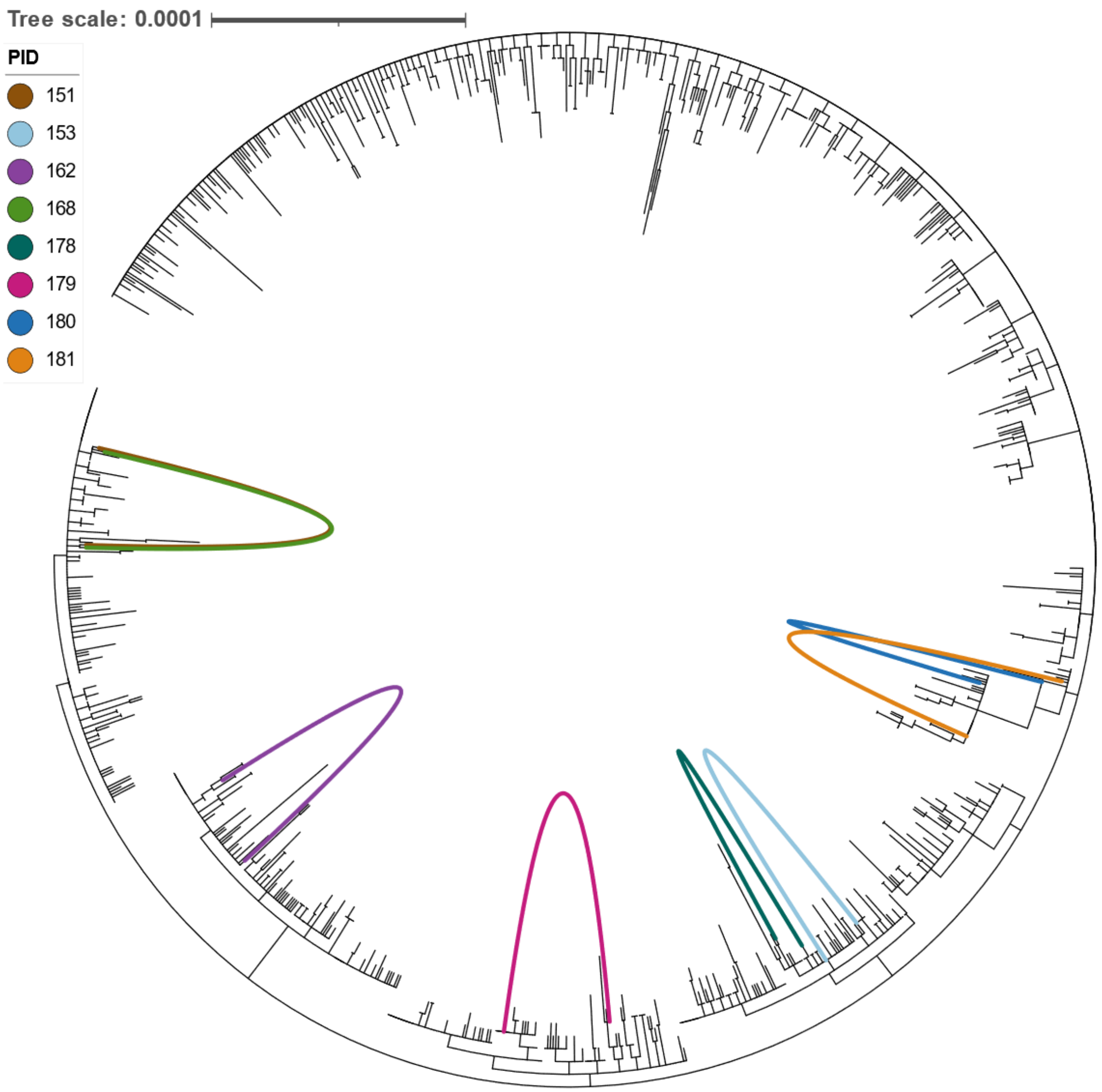


**Supplementary Figure 7**. **Infection with multiple MPXV strains in the NYC individuals with sequences which are distantly related in the phylogeny**. This NYC MPXV tree was inferred from 1,114 NYC sequences by masking the region with low depth of coverage. Links in the figure were used to connect the sequences from the same individual that had been categorized to distantly related using their phylogenetic placement. Sequences from Individual ID 151 (brown color) and Individual ID 168 (green color) were very closely placed on the phylogeny. The joining lines (brown and green) appeared to be overlaying each other.

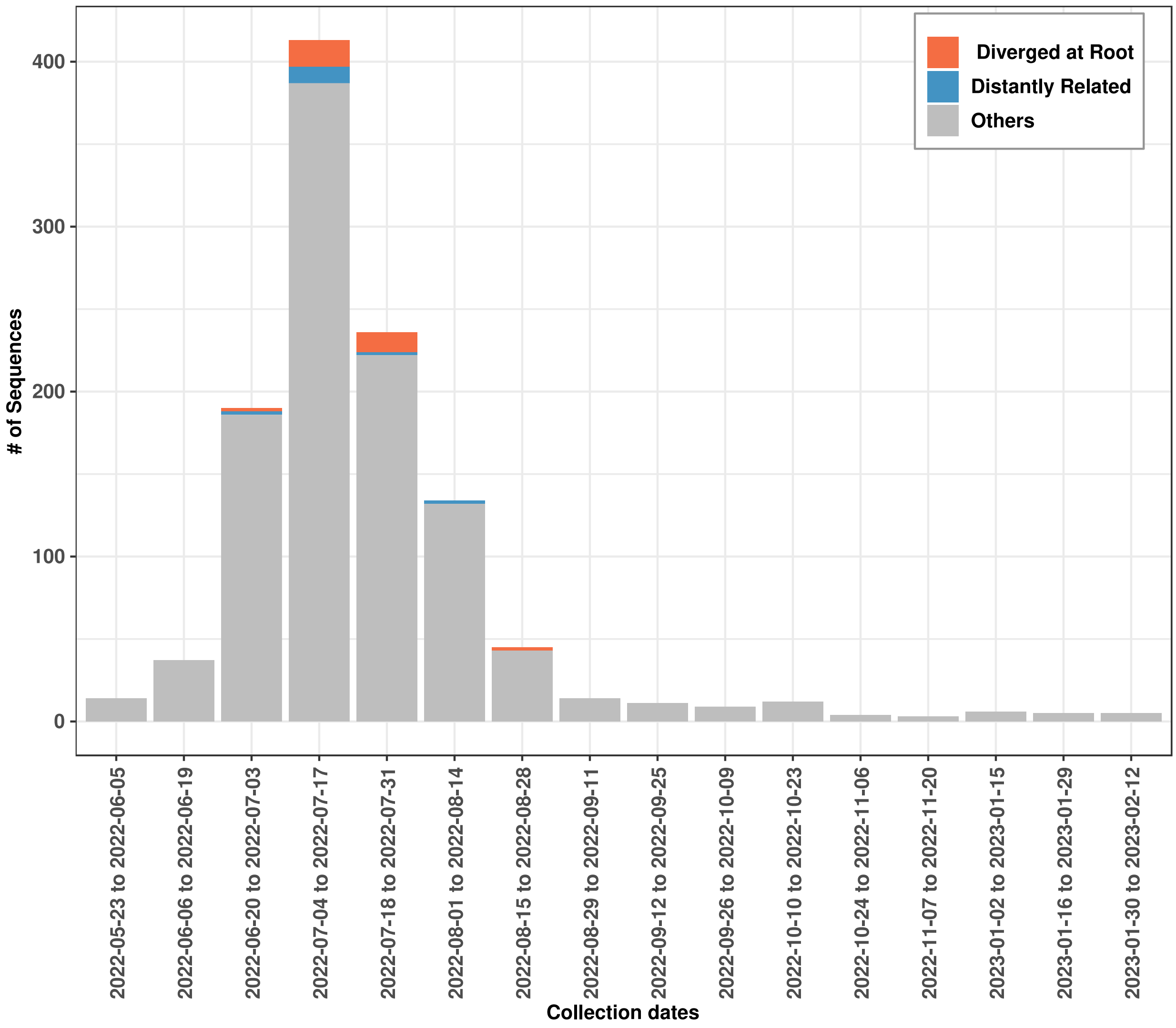


**Supplementary Figure 8**. **Temporal distribution of 1,138 NYC genomes sequenced from the individuals with and without infection with multiple MPXV variants**. Sequencing was done at the NYC Public Health Lab (PHL). The y-axis denotes the total number of sequences. The x-axis denotes the biweekly collection dates. Sequences were from the individuals with a confirmed case of infection with multiple MPXV variants (Diverged at Root), probable case of multiple infection (Distantly Related) and without multiple infection (Others). Most genomes sequenced from individuals with confirmed and probable cases of infection with multiple MPXV variants were collected from July 4$^{th}$ – July 31, 2022.

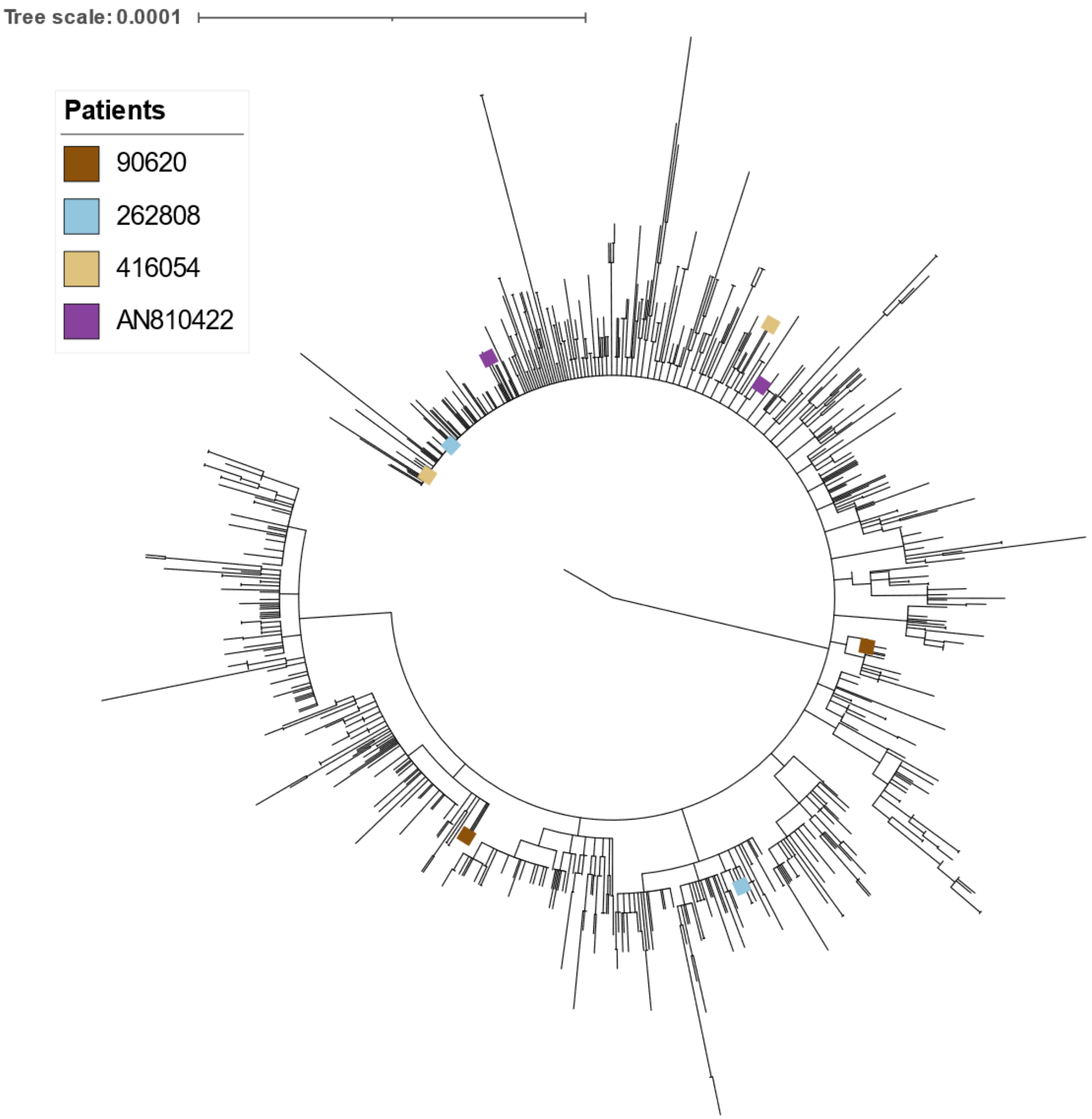


**Supplementary Figure 9**. **Intra-host phylogeny between sequences sampled from different lesions in the same individual**. Patients with larger phylogenetic distances are visualized in **Figure 5.** A phylogenetic distance greater than 0.000021 was annotated with colored tips and plotted on the NYC only phylogeny. One individual's sequences (90620/brown) resulted in non-monophyletic placements of sequences.

### Supplementary Tables

**Supplementary Table 1**. B lineage specific mutations in the global (Including NYC PHL) MPXV dataset.

| Position | Mutation | Effect | Gene | APOBEC |
|---|---|---|---|---|
| 1262 | S105L | NON_SYNONYMOUS_CODING | OPG001 | APOBEC |
| 2591 | S54F | NON_SYNONYMOUS_CODING | OPG002 | APOBEC |
| 3111 | I499 | SYNONYMOUS_CODING | OPG003 | APOBEC |
| 3522 | I362 | SYNONYMOUS_CODING | OPG003 | APOBEC |
| 3818 | D264N | NON_SYNONYMOUS_CODING | OPG003 | APOBEC |
| 7771 | I64 | SYNONYMOUS_CODING | OPG019 | APOBEC |
| 14000 | A423D | NON_SYNONYMOUS_CODING | OPG025 | NON-APOBEC |
| 15428 | *NA* | INTERGENIC | *NA* | APOBEC |
| 21723 | F237 | SYNONYMOUS_CODING | OPG037 | APOBEC |
| 23564 | S185 | SYNONYMOUS_CODING | OPG039 | APOBEC |
| 25661 | *NA* | INTERGENIC | *NA* | APOBEC |
| 30367 | L276 | SYNONYMOUS_CODING | OPG047 | APOBEC |
| 31053 | R48C | NON_SYNONYMOUS_CODING | OPG047 | APOBEC |
| 34459 | P78S | NON_SYNONYMOUS_CODING | OPG053 | APOBEC |
| 37202 | F611 | SYNONYMOUS_CODING | OPG056 | APOBEC |
| 38360 | I225 | SYNONYMOUS_CODING | OPG056 | APOBEC |
| 38662 | E125K | NON_SYNONYMOUS_CODING | OPG056 | APOBEC |
| 39119 | E359 | SYNONYMOUS_CODING | OPG057 | APOBEC |
| 39139 | E353K | NON_SYNONYMOUS_CODING | OPG057 | APOBEC |
| 52885 | V518 | SYNONYMOUS_CODING | OPG071 | APOBEC |
| 54117 | L108F | NON_SYNONYMOUS_CODING | OPG071 | APOBEC |
| 54635 | D56N | NON_SYNONYMOUS_CODING | OPG072 | APOBEC |
| 64297 | I140 | SYNONYMOUS_CODING | OPG083 | APOBEC |
| 72362 | D196N | NON_SYNONYMOUS_CODING | OPG092 | APOBEC |
| 73066 | S30L | NON_SYNONYMOUS_CODING | OPG093 | APOBEC |
| 73239 | D88N | NON_SYNONYMOUS_CODING | OPG093 | APOBEC |
| 74205 | M142I | NON_SYNONYMOUS_CODING | OPG094 | APOBEC |
| 77383 | E162K | NON_SYNONYMOUS_CODING | OPG098 | APOBEC |
| 81275 | K50 | SYNONYMOUS_CODING | OPG105 | APOBEC |
| 82373 | F416 | SYNONYMOUS_CODING | OPG105 | APOBEC |
| 82451 | T442 | SYNONYMOUS_CODING | OPG105 | APOBEC |
| 83326 | S734L | NON_SYNONYMOUS_CODING | OPG105 | APOBEC |

| | | | | |
|---|---|---|---|---|
| 84587 | F1154 | SYNONYMOUS_CODING | OPG105 | APOBEC |
| 87230 | H740Y | NON_SYNONYMOUS_CODING | OPG109 | APOBEC |
| 87297 | F717 | SYNONYMOUS_CODING | OPG109 | APOBEC |
| 91728 | *NA* | INTERGENIC | *NA* | APOBEC |
| 95034 | V125 | SYNONYMOUS_CODING | OPG115 | NON-APOBEC |
| 119296 | D98N | NON_SYNONYMOUS_CODING | OPG136 | APOBEC |
| 121320 | A17T | NON_SYNONYMOUS_CODING | OPG139 | NON-APOBEC |
| 124130 | E62K | NON_SYNONYMOUS_CODING | OPG145 | APOBEC |
| 124674 | R243Q | NON_SYNONYMOUS_CODING | OPG145 | APOBEC |
| 125249 | E435K | NON_SYNONYMOUS_CODING | OPG145 | APOBEC |
| 128698 | S307L | NON_SYNONYMOUS_CODING | OPG150 | APOBEC |
| 133175 | *NA* | INTERGENIC | *NA* | NON-APOBEC |
| 148412 | I328 | SYNONYMOUS_CODING | OPG174 | APOBEC |
| 150469 | H221Y | NON_SYNONYMOUS_CODING | OPG176 | APOBEC |
| 151461 | *NA* | INTERGENIC | *NA* | NON-APOBEC |
| 155795 | *NA* | INTERGENIC | *NA* | APOBEC |
| 162243 | L216 | SYNONYMOUS_CODING | OPG188 | APOBEC |
| 162331 | L246 | SYNONYMOUS_CODING | OPG188 | APOBEC |
| 164832 | L500 | SYNONYMOUS_CODING | OPG189 | APOBEC |
| 168120 | L263F | NON_SYNONYMOUS_CODING | OPG193 | APOBEC |
| 170262 | R75 | SYNONYMOUS_CODING | OPG198 | APOBEC |
| 178133 | *NA* | INTERGENIC | *NA* | APOBEC |
| 181980 | D209N | NON_SYNONYMOUS_CODING | OPG210 | APOBEC |
| 183519 | P722S | NON_SYNONYMOUS_CODING | OPG210 | APOBEC |
| 186578 | M1741I | NON_SYNONYMOUS_CODING | OPG210 | APOBEC |
| 187428 | *NA* | INTERGENIC | *NA* | APOBEC |

**Supplementary Table 2**. Accession #, lineage assignment and analysis results of groups epidemiologically linked cases in NYC MPXV dataset:

- See file Supplementary_Table_2_Epilinked_results.xlsx

**Supplementary Table 3**. MPXV genomes sequenced at the NYC public health laboratory (NCBI accession and metadata-Ct, genome coverage, Lineage, collection date):

- See file Supplementary_Table_3_&_4_MPXV_genome_sequences.xlsx: NYC_PHL

**Supplementary Table 4**. Accession numbers, collection dates, geography, and lineages in global MPXV dataset:

- See file Supplementary_Table_3_&_4_MPXV_genome_sequences.xlsx: Global.

**Supplementary Table 5.** Regions masked due to long stretches of low depth in >10% sequences and the 3' ITR region.

| Masked_Region | reference | posStart | posEnd | mask_length | Note |
|---|---|---|---|---|---|
| 1 | NC_063383.1 | 1 | 770 | 770 | |
| 2 | NC_063383.1 | 12860 | 13064 | 205 | |
| 3 | NC_063383.1 | 31058 | 31715 | 658 | |
| 4 | NC_063383.1 | 70475 | 71209 | 735 | |
| 5 | NC_063383.1 | 71932 | 72341 | 410 | |
| 6 | NC_063383.1 | 89636 | 90344 | 709 | |
| 7 | NC_063383.1 | 99192 | 99341 | 150 | |
| 8 | NC_063383.1 | 100440 | 101060 | 621 | |
| 9 | NC_063383.1 | 113539 | 113954 | 416 | |
| 10 | NC_063383.1 | 142485 | 143105 | 621 | |
| 11 | NC_063383.1 | 196439 | 197209 | 771 | |
| 12 | NC_063383.1 | 190788 | 197209 | 797211 | 3' ITR region |
